# A universal power law optimizes energy and representation fidelity in visual adaptation

**DOI:** 10.1101/2025.03.20.643406

**Authors:** Matteo Mariani, S. Amin Moosavi, Dario L. Ringach, Mario Dipoppa

## Abstract

Sensory systems continuously adapt their responses based on the probability of encountering a given stimulus. In the mouse primary visual cortex (V1), the population response magnitude is a power law of the stimulus probability in the environment. For a given stimulus type (e.g., oriented gratings), the power law’s exponent is invariant to changes in statistical environments, enabling predictions of population responses to new environments. Here, we aim to provide a normative explanation for the power law behavior. We develop an efficient coding model where neurons adjust their firing rates through optimization of a weighted objective, hypothesizing that the neural population adapts to enhance stimulus detection and discrimination while reducing overall neural activity. We show that a model balancing representational fidelity and energy efficiency matches the power law observed experimentally for a wide range of parameters, while models of adaptation with alternative coding objectives and resource constraints are unable to reproduce this empirical observation. Furthermore, we account for the invariance of the power law’s exponent across environmental changes by linking it to the dependence of tuning curve modulation on stimulus probability. Finally, we explain how variations in the exponent with different stimulus types (e.g., natural stimuli) result from changes in the minimal distances between neural representations, in agreement with experimental findings. We conclude that a universal power law of adaptation can be explained as a trade-off between representation fidelity and energy cost.

## Introduction

Adapting to new environments is a hallmark of intelligent systems. Sensory systems continuously adapt their representation of stimuli based on environment statistics [1–7]. In the visual system, adaptation occurs across several areas of the brain, from the retina [8, 9] to the primary visual cortex (V1) [1, 2, 10–12] and in higher visual areas, such as V2 [11], V4 [13] and MT [14], involving different time scales, from milliseconds to seconds [6, 12, 15, 16] and different stimulus features, such as stimulus contrast [8, 10, 11, 16] and orientation [1, 17–19]. Adaptation also impacts our sensory experience, generating visual illusions in perception of orientations [20, 21] and higher visual features, such as faces [22].

Several theoretical frameworks, not always mutually exclusive, have been proposed to explain the mechanisms and functional benefits of adaptation [23]. These paradigms include redundancy reduction [24–26], homeostatic drive of responses [2, 27], predictive coding [28–31], surprise salience [32, 33], Bayesian inference [34, 35], and efficient coding [36–39]. According to the efficient coding paradigm, neurons maximize the information about the environment they represent, subject to constraints, such as sparseness, limited energy, limited dynamic range, finite internal noise [40–43]. Efficient coding has been particularly successful in explaining how neural populations adapt to the statistics of the environment [39, 44].

Characterizing adaptation-induced changes in neural populations is crucial to fully understand how representations are transformed during adaptation [39, 45]. Experimental advances have enabled us to study adaptation in populations of hundreds of neurons [1, 10, 39, 46]. A remarkable finding in one of these large-scale recording studies is that the population response magnitude to a given stimulus is proportional to a power of its occurrence probability [1]. In that study, during a set time block, stimuli were sampled according to different probability distributions, which defined a statistical sensory environment for that time block. For a given stimulus type, the power law exponent was invariant across probability distributions of stimuli (Figure 1d).

**Figure 1.**
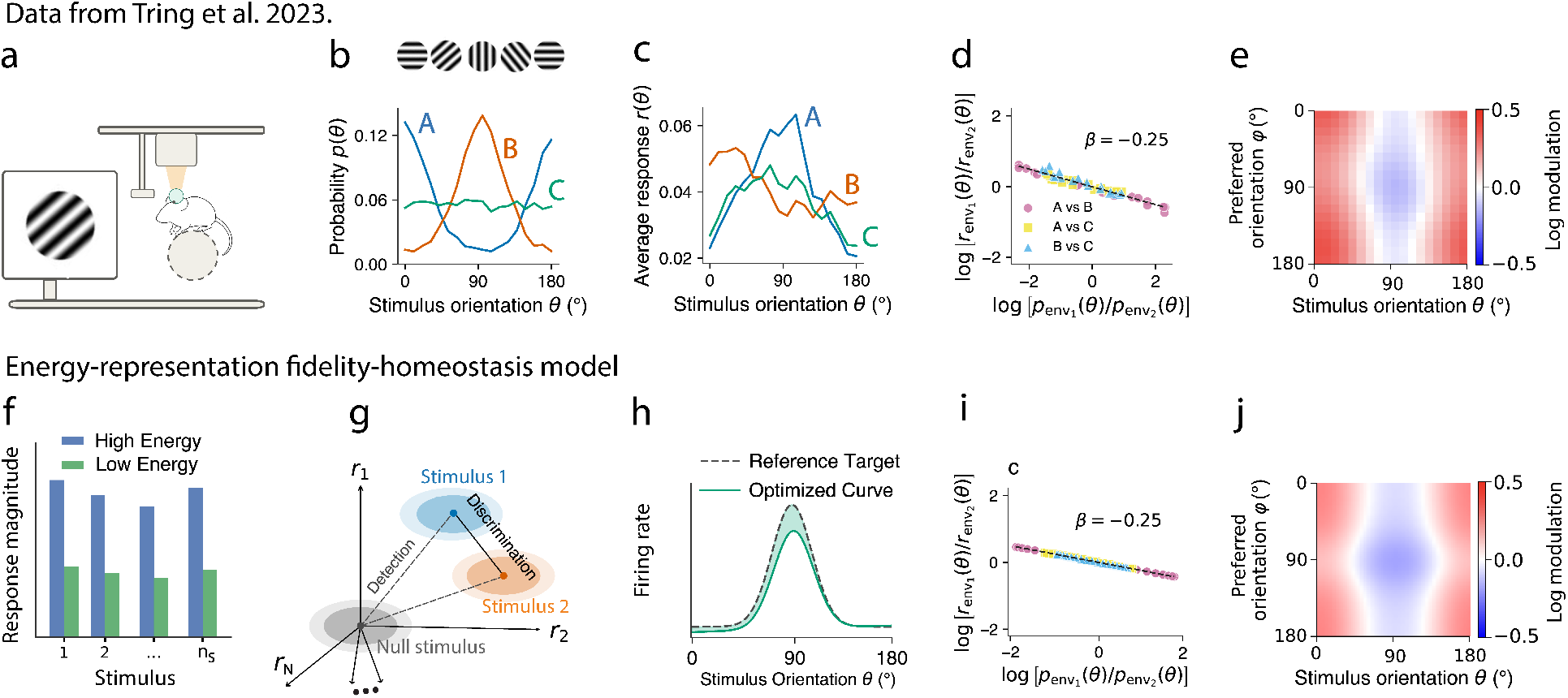
An efficient coding model to explain the universal power law of visual adaptation. a) Schematic of the experimental setup in [1]. b) Three environments of oriented gratings from [1]. Each environment is characterized by a probability distribution of the grating orientations. c) The average population responses to each of the environments in (b). The responses are paired with the environments by color. d) Ratio of average population responses vs ratio of probabilities, for the three environments in (b). Data re-analyzed from [1]. Each colored marker represents a unique pair of environments. e) Adaptation induced modulation of the neural responses in the data [1], computed as the ratio between responses in biased vs uniform environment. f-h) We pose that adaptation is driven by four objectives, the push of saving energy (f), of reducing discrimination and detection errors (g), and a homeostatic constraint on the responses (h). i) Same as d), but obtained from the model. j) Same as (e), but obtained from the model.

In the present study, we show that a normative efficient coding model reproduces the power-law of adaptation across a broad range of parameters. We find that the invariant power law scaling emerges from a fundamental trade-off between energetic cost and representational fidelity, quantified by the ability to detect and discriminate visual stimuli. We compared the result with alternative models of adaptation and showed that they cannot capture the key features of the empirical data. We then fitted the model to neural responses, and further showed that it provides predictions for how neural responses adapt across environments of stimuli. Importantly, we show that change in stimulus type alters representation fidelity, which in turn changes the exponent of the resulting power law due to a competition between two discrimination mechanisms, a gain difference term and an orthogonality term. Together, these results suggest that the invariant power law of adaptation emerges from the competing demands of energetic efficiency, stimulus detection and discrimination enhancement.

## Results

Two fundamental properties of adaptation in mouse primary visual cortex emerged in a prior experimental study (Figure 1a-e): 1) the ratio of the average population responses is a power law of the ratio of stimulus probabilities for two different environments, and 2) the power law exponent is invariant to changes in the probability distributions [1]. To mathematically specify these two properties, we first define an environment. For a general set of *n*_*S*_ stimuli 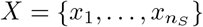, an environment *e* is defined by the probability distribution *p*_*e*_ : *X* → [0, 1] with which stimuli are presented. For example, for a set of oriented gratings, *p*_*e*_(*θ*) is the probability in an environment *e* that a grating with orientation *θ* ∈ [0, *π*] is presented (Figure 1b). When a stimulus *θ* is presented to a subject, it triggers a set of responses in the neural population (Figure 1c). Due to adaptation, the responses do not depend solely on the stimulus, but also on the environment. To quantify such dependence, we define, for a given stimulus *x* and environment *e*, both the response vector **r**_*e*_(*θ*) = {*r*_*e*,1_(*θ*), *r*_*e*,2_(*θ*), …, *r*_*e,N*_ (*θ*)}, that is the vector of the firing rates of each of the *N* neurons, and the average population response, 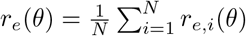.

Following the above definition of environment and average population response, we can mathematically characterize the first property of adaptation observed in [1]: given two environments *env*_1_ and *env*_2_ (Figure 1b), the ratio of the average population responses as a function of the average environment probabilities is described by the following power law (Figure 1d):

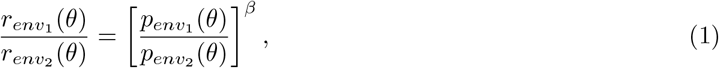

We can then mathematically characterize the second key property of adaptation observed in [1]: the exponent *β* is approximately invariant to the choices of environments.

We introduced an efficient coding model to explain which trade-off between coding objectives and resource constraints underlies the emergence of the invariant power law. Let us consider a homogeneous set of reference tuning curves *r*_0_ (*φ, θ*), namely tuning curves with the same functional form and amplitude and equally-spaced preferred orientations, where *φ* indicates each neuron’s preferred orientation and *θ* is the stimulus orientation, both parametrized between 0 and *π*. Tuning curves of different neurons differ only by translation, reflecting their distinct preferred orientations. To find the adapted tuning curves *r*_*e*_(*φ, θ*) in a given environment *e*, the model numerically computes a function *a*_*e*_(*φ, θ*) modulating the reference tuning curves *r*_0_ (*φ, θ*) as follows: *r*_*e*_(*φ, θ*) = *r*_0_(*φ, θ*)*a*_*e*_(*φ, θ*). Such modulation emerges from a weighted-objective optimization that we hypothesized captures the principles of sensory adaptation. The adapted neural responses thus emerge from the minimization of a cost function that includes: an energy cost (Figure 1f), reflecting the need to reduce metabolic expenditure; terms that promote representation fidelity of stimuli (Figure 1g), ensuring that sensory responses remain informative within a given environment; homeostatic drives that enforce physiological plausibility in the resulting tuning of the neural responses (Figure 1h). The adapted tuning curves minimize thus the following total cost:

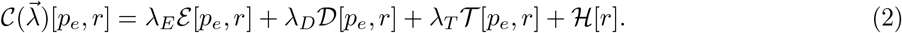

Each of these terms contributes to shaping the optimal modulation of the tuning curves. The term ℰ [*p*_*e*_, *r*] is the expected *energy cost* (i.e., the metabolic consumption of firing action potentials), over the probability *p*_*e*_(*θ*), of encoding the stimuli in the environment *e*. The energy cost penalizes a high average population response.The next two terms take into account representation fidelity. The term *D*[*p*_*e*_, *r*] is the expected *discrimination error* that we defined by means of the Mahalanobis distance of the response vectors [47] (Methods), over all pairs of stimuli (*θ*_*i*_, *θ*_*j*_) averaged over the probability *p*_*e*_(*θ*_*i*_) · *p*_*e*_(*θ*_*j*_). The discrimination error penalizes small distances between response vectors encoding different stimuli relative to the noise. The term *T* [*p*_*e*_, *r*] represents the expected *detection error* across all stimuli, quantifying the ability to distinguish the presence of a stimulus from noise. The *homeostasis constraint* ℋ [*r*] induces the adapted tuning curves to be similar to the reference tuning curves *r*_0_ (*φ, θ*). The weights 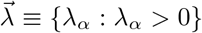, with *α* ∈ {*E, D, T, H*}, are hyperparameters of the model and represent the relative importance of each term in the optimization. One of the weights would be redundant so we set *λ*_*H*_ = 1. Finding the adapted tuning curves to an environment *e* for a set of weights 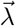 reduces to finding the modulation *a*_*e*_(*φ, θ*) such that the tuning curves *r*_*e*_(*φ, θ*) = *r*_0_(*φ, θ*)*a*_*e*_(*φ, θ*) are the minimizers of the functional *C*, namely 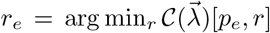. The optimization was performed iteratively by gradient descent subject to the following constraint: *a*_*e*_(*φ, θ*) = *a*_*e*,1_(*φ*)*a*_*e*,2_(*θ*), where *a*_*e*_, *i* : [0, *π*] → ℝ^+^ for *i* = 1, 2 which are respectively the neural and the stimulus modulations.

### The power law of adaptation emerges from an efficient coding model

A power law emerged as the solution of the efficient coding model. We considered the three environments of [1] in Figure 1b: one defined by a uniform probability distribution and the other two by von Mises probability distributions. For each environment *e* ∈ {*A, B, C*} we used our model to find the predicted tuning curves *r*_*e*_(*φ, θ*). We next computed the average population responses *r*_*e*_(*θ*) for each environment by averaging over the neurons’ preferred orientation. The ratio of the average population responses as a function of the ratio of probabilities followed a power law (Figure 1i) as in the experimental data [1]. Note that, because in the model the reference tuning curves are homogeneous, we obtain a power law for the average population response as a function of the environment probability for any environment *e*, namely *r*_*e*_(*θ*) ∝ *p*_*e*_(*θ*)^*β*^, without the need to consider the ratios of environment pairs. However, in the experiments, it is necessary to take such ratios to obtain a power law, because the tuning curves are inhomogeneous in all environments, including the uniform one. We first present the results for the case of homogeneous reference tuning curves, to focus on the effect of short-term adaptation, abstracting away from inhomogeneities. Later, we will also show that the efficient coding model reproduces the power law in the case of inhomogeneous tuning curves (Figure 5).

The modulation of tuning curves predicted by the model matches the modulation that we found in the data. We performed in fact a novel analysis of the experimental data in [1], in which we computed the adaptation-induced modulation by evaluating the ratio of tuning curves in a von Mises environment to those in a uniform environment, averaged across recording sessions. The modulation exhibited stronger dependence across stimuli than across neurons (Figure 1e). By performing the same analyses for the adapted tuning curves, as predicted by the model, we found a matching result for the modulation (Figure 1j). The accurate prediction of tuning curves modulation demonstrates that the model captures not only global, population-level effects, but also local changes due to adaptation.

The model’s results are robust when changing the noise distribution. The model assumes Gaussian, independent, unit-variance noise in neural responses. We studied whether the power law could also arise under different noise statistics. We thus considered Poisson noise by modeling each neuron as an independent generator of Poisson spike trains. The model predicts adapted tuning curves that reproduce the power law also when the noise is Poisson-distributed (Suppl. Figure S.1a-f).

All four terms of the cost function of the model are necessary to reproduce the key set of experimental findings in [1]. The energy cost, detection error, and homeostasis constraint were necessary to obtain a power law with a negative exponent. To see this, we considered a von Mises environment (Suppl. Figure S.2a) and we set to zero, one at a time, each of the weights *λ*_*α*_ of the cost function. Removing the energy cost led to an average population response that increases with the probability (Figure S.2b). Removing the homeostasis constraint flattens the adapted average population response to almost a constant (Suppl. Figure S.2b). The presence of the homeostasis constraint maintains the average population response closer to the reference one, forcing the average population response to be a non-constant function of the probability. Removing the detection error breaks the power law as well (Suppl. Figure S.2b). The discrimination error was necessary to capture the recorded neuron-dependent modulation of the responses [1], which cannot be reproduced if we remove the discrimination error (Suppl. Figure S.2c). This arises from the mathematical structure of the cost terms: the discrimination cost is the only cost to depend on the relative direction of the neural response vectors (Methods). Consequently, including the discrimination error in the weighted objective cost modulates responses in a neuron-dependent manner. The discrimination error is shaped by the optimization process even if it is not explicitly included in the total cost function (Suppl. Figure S.3). Specifically, changes in detection affect changes in discrimination (Suppl. Figure S.3b-c) because increasing the magnitude of the responses improves both detection and discrimination.

### The power law exponent is independent of the environment and reflects the relative magnitude of the model’s weights

The model predicted an invariant power law to changes in the statistical environment for a wide range of parameters. We studied the output of the model in several environments (Figure 2a) compatible with the experimental ones [1], for fixed values of the weights *λ*_*α*_. We sampled pairs of environments and observed that the relation between the ratio of probabilities and the ratio of average population responses was approximately invariant across pairs of environments, allowing us to fit all the points of the model with one power law (Figure 2b-c). We then allowed the detection weight *λ*_*T*_ and the energy weight *λ*_*E*_ to vary over 6 orders of magnitude and we computed the average *R*^2^ for the power law fit (Figure 2d) and the average invariance score *ω* (Figure 2e). The model showed both a large coefficient of determination *R*^2^ and a large invariance *ω* for a wide range of the parameters. The power law exponent depended on the relative magnitude of the model’s weights (Figure 2f), showing that the exponent value is a signature of a trade-off between conflicting constraints: energy efficiency and representation fidelity.

**Figure 2.**
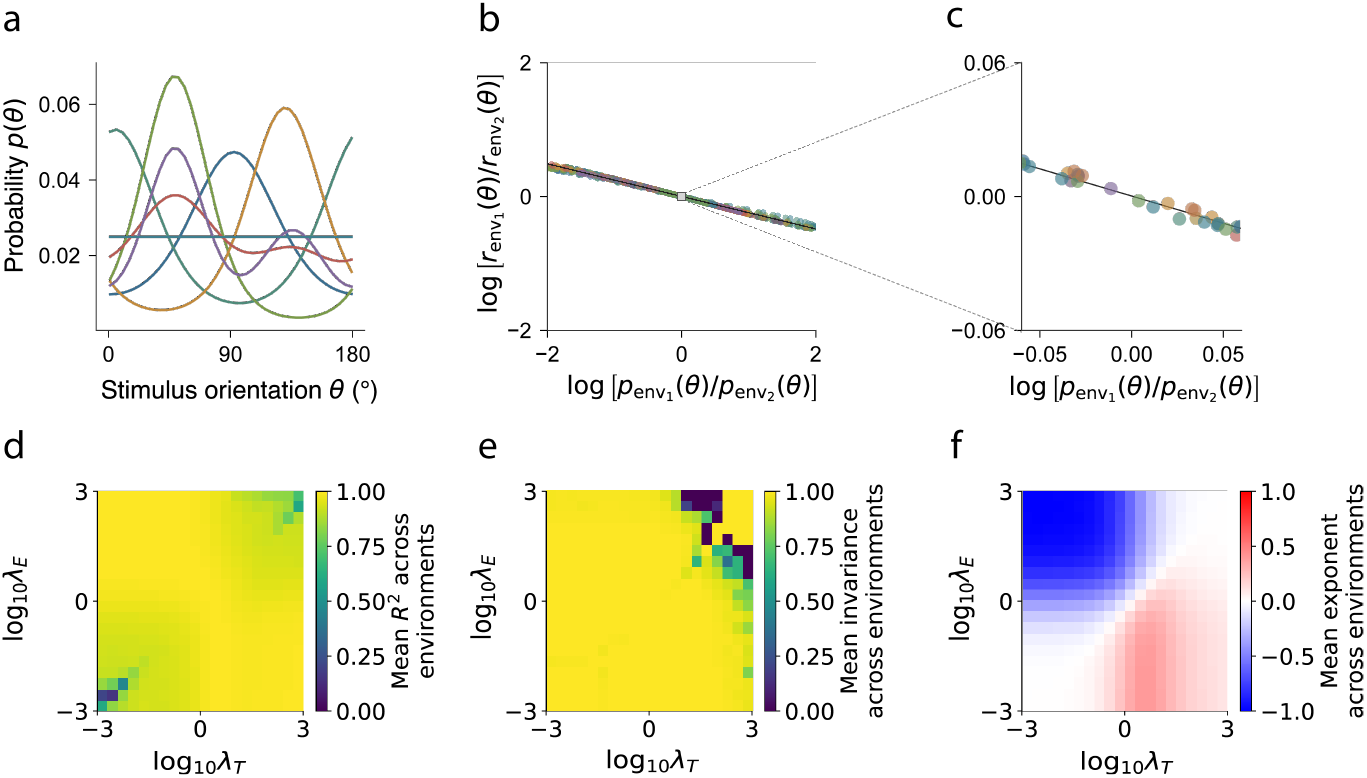
The model reproduces an environment-invariant power law for a wide set of parameters. a) Set of example environments defined by different probability distributions of the stimulus orientation. b) Colored circles: ratios of response rates as a function of the ratio of probabilities from different pairs of environments in (a); solid line: power law fit. Response rates were computed with a fixed weight vector *λ*_*α*_. c) Magnified view around the origin to appreciate negligible variations across pairs of environments. d-f) Average over the environments of: the quality *R*^2^ of the power law fit (d), the invariance score *ω* (e), the power law exponent as functions of the energy and detection weights. The homeostasis weight was fixed to *λ*_*H*_ = 1 and the discrimination weight to *λ*_*D*_ = 0.3.

Responses in mouse V1 are known to be amplified during running [48, 49], raising the question of whether locomotion also modulates the exponent. We found that locomotion changes the tradeoff between energy and representation fidelity by increasing the magnitude of responses, without however affecting the power law exponent. We compared in fact the power law in two locomotion states: running and rest. We separated trials by locomotion state (running vs. rest) in data from [1], and measured the exponent in each condition. We found no significant change in the exponent across experiments between running and rest states (Figure S.4a, Wilcoxon test: *p* = 0.92), while the overall response amplitude is significantly increased during running (Figure S.4b). The power-law exponent remains unchanged while neural responses are amplified during running if responses follow the relation *r*(*θ*) = *c*_*s*_ *p*(*θ*)^*β*^, where *c*_*s*_ depends on the locomotion state *s* ∈ {rest, run}. The exponent *β* remains invariant across states. Taking the logarithm yields log *r*(*θ*) = *β* log *p*(*θ*) + log *c*_*s*_, indicating that locomotion shifts the power law vertically in log-log coordinates without altering its slope. The model accounts for this effect. Adjusting the relative weights of energy and detection in a simplified version of the model produces two parallel lines when plotting the average population response as a function of stimulus probability (Figure S.4c).

### Alternative models of adaptation cannot capture empirical features of the environment-invariant power law

We studied alternative prominent normative models of adaptation, which are grounded on different trade-offs between coding objectives and resource constraints, and observed that, among the models considered, only a model with our weighted objective function successfully reproduced all the key features of the data. All the alternative models considered aim to find the functional advantages of adaptation, but differ in the specific benefit achieved by the visual system. For example, homeostasis of different response statistics has been considered a driver of adaptation. While homeostasis of average responses [50], corresponding to the first moments of the response distribution, has been ruled out in mouse V1 by [1], whether homeostasis of the second moments, namely pairwise product and covariance of responses [2] can account for the experimentally observed power law remains an open question. We considered two variants of the models enforcing respectively the covariance and the product homeostasis (Figure 3(ii-iii)). For each of the models, we performed a grid search over their hyperparameters and selected the one that was closest to the data, namely that exhibited an invariant power law (large *R*^2^ and *ω*) with exponent close to the data (Methods). We first studied whether a power law emerged for each of these models for a fixed pair of environments. We observed that the covariance homeostasis model does not exhibit a power law (Figure 3aii), while the product homeostasis does (Figure 3aiii). To assess invariance of the power law, we considered multiple environments. The product homeostasis model exhibits an invariant power law, which, however, does not match the exponent of the data (Figure 3biii). Then we assessed the robustness of the models considered across weights of the cost function, by representing the density of parameters in the grid search as a function of the quality of the power law fit *R*^2^ and the invariance score *ω*. The product homeostasis model exhibits a high density of parameters around the best parameter chosen (Figure 3ciii), while the best model of covariance homeostasis is far from the high density zone (Figure 3cii). Finally, we compared the modulation to a biased von Mises environment predicted by each model’s best hyperparameter selected. Both covariance and product homeostasis models predict a larger contribution of neural modulation (Figure 3d(ii-iii)) than the one seen in the data (Figure 1e).

**Figure 3.**
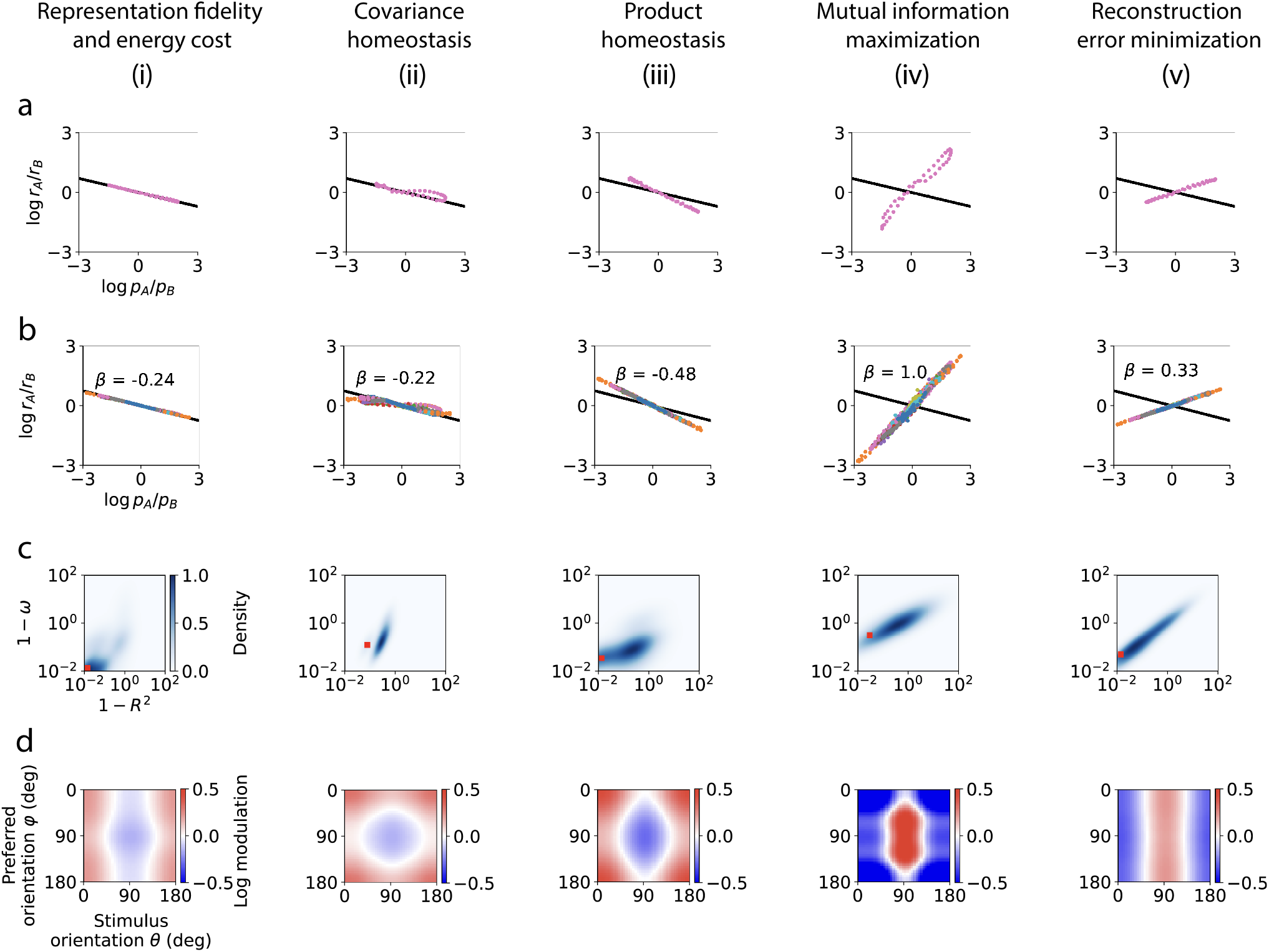
Alternative models of adaptation cannot reproduce the main features of the data. i-v) Five models of adaptation considered, in order: i) representation fidelity and energy cost model, ii) covariance homeostasis model, iii) product homeostasis model, iv) mutual information maximization model, v) reconstruction error minimization model. Each model was evaluated according to four assessment criteria corresponding to rows a-d). a) Power law existence for the best hyperparameter set of each of the five models. The dots represent the model’s output, while the black line is the power law with the exponent of the data in [1]; b) Power law of each model for its best set of hyperparameters. Differently colored dots represent different pairs of environments. The black solid line represents the power law with the exponent of the data. c) The density of each model’s hyperparameters as a function of their power law *R*^2^ ∈ (−∞, 1] and invariance *ω* ∈ (−∞, 1]. The axes represent 1 − *R*^2^ and 1 − *ω* so that the best possible value is the origin. The red square is the hyperparameter set chosen for each model and used in rows (a,b,d). d) Tuning curves modulation predicted by each model to a biased von Mises environment. For comparison, the modulation of the data is in Figure 1e.

In alternative to homeostasis, the maximization of local representation fidelity with limited coding constraint has been considered as a possible driver of adaptation [37, 38, 51]. While our model defines fidelity of representation as a weighted sum between detection and discrimination, in turn defined through Mahalanobis distance, the fidelity of representations has also been defined by means of mutual information [47], usually in a local approximate form based on Fisher information [37, 38, 47]. We thus analyzed a variant of a model that maximizes the mutual information under a constraint of limited coding capacity [36, 38]. A generalization of this model consists in minimizing the reconstruction error (estimated again using a power of the Fisher information) [51]. As in the case of the homeostasis models, we performed a grid search over the hyperparameters and picked the one closest to the data. Both the mutual information and the reconstruction error models exhibited an approximate power law (Figure 3a(iv,v). We observed that the power law was also invariant for both models. However, the exponent did not match the data (Figure 3b(iv,v)). The exponent was in fact positive, and this can be derived analytically when the modulation is only stimulus dependent (Supplementary information). We then assessed the robustness of the efficient coding models: both the mutual information and the reconstruction error models exhibited high density of parameters around the best parameter chosen (Figure 3c(iv,v)). The analysis of the modulation showed that the mutual information and the reconstruction error models have a modulation with inverted sign compared to the experimental data (Figure 3d(iv-v)). This is a direct consequence of their positive power law exponent.

A model optimizing energy and local reconstruction error can provide a negative invariant power law, but with an unrealistic modulation of tuning curves. To show this, we considered the efficient coding model of [37, 52], that optimizes a monotonic function of the Fisher information, constrained by limited energy, by allowing a redistribution of neurons’ preferred orientation and a change in tuning sharpness (Suppl. Figure S.5a-c). The model exhibited a negative invariant power law with an exponent compatible with the data (Suppl. Figure S.5d). However, the change in neural density of preferred orientation gives rise to a modulation (Suppl. Figure S.5e) that is substantially different from the results in [1], showing that this model cannot describe the experimentally observed adaptation effects.

Among the models considered above, the one combining energy, discrimination, detection and homeostasis was thus the only to match the main features of the data of [1], since it exhibited a power law (Figure 3ai) that was environment-invariant and with an exponent close to the data (Figure 3bi), for a parameter set that was picked in a high-density region of parameters (Figure 3ci), and with a modulation very close to the data (Figure 3di).

### The emergence of the environment-invariant power law is explained analytically by a simplified model

Although all the terms in the cost function are necessary to reproduce the key experimental findings reported in [1] (Figure S.2), only energy, homeostasis, and detection are sufficient to reproduce the power law alone. To better understand the specific contributions of the energy cost, detection error, and homeostasis constraint, we analyzed a simplified model including only these three terms. We found that the power law is approximate (as in Figure 1i), yet there exists a region of the parameter space in which the approximation error is minimal (Figure 4a) and the exponent varies smoothly from −1 to 1 (Figure 4b-c).

**Figure 4.**
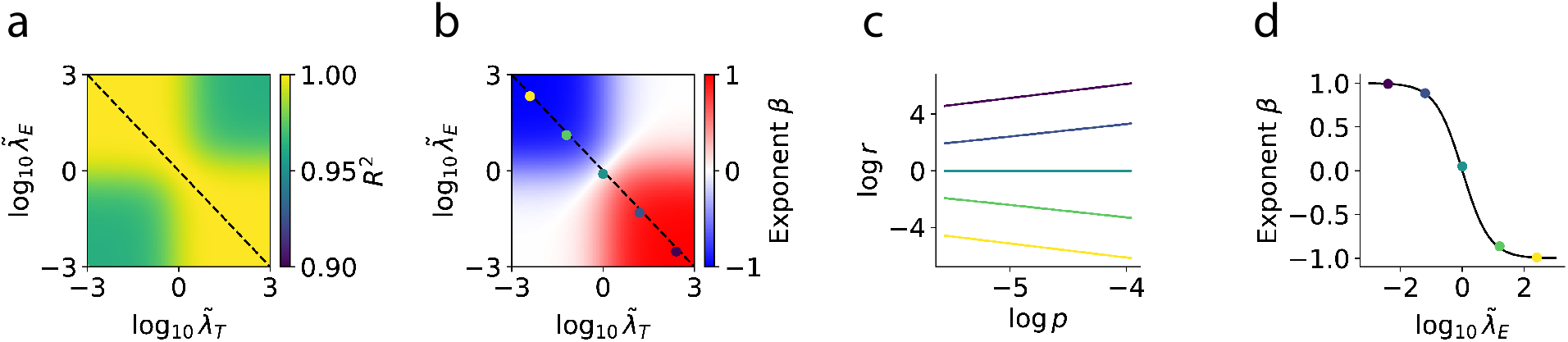
An analytic approach finds an approximate power law for the response rate. a) The *R*^2^ of the power law approximation of the modulation, shown as a function of the detection and energy weights rescaled by the homeostasis weights: 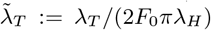 and 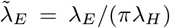. *Dashed* : the line of parameters which give the maximal *R*^2^: it is the line subject to the constraint 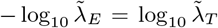. b) The exponent of the best power law fitting the modulation. *Dashed* : same as (a). Colored dots: points on the line of best approximation of power law. c) Each colored line corresponds to a different color in the previous panel. d) Colored dots: the exponent corresponding to each point marked in panel (b). Solid line: exponent predicted by a Taylor expansion (Supplementary Information).

The simplified model provides insight into the mathematical origin of the environment-invariant power law. This invariance arises because, for every stimulus orientation *θ*, the average population response *r*_*env*_(*θ*) depends exclusively on the local stimulus probability *p*_*env*_(*θ*) and not on the functional form of the probability distribution. By calculus of variations, indeed, we found that there exists a function *f*_*λ*_ such that:

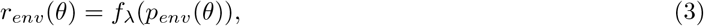

where:

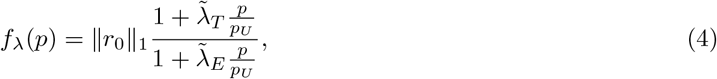

where *p*_*U*_ denotes the probability under the uniform environment, 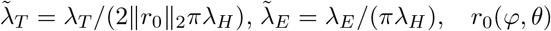 represents the reference tuning curves, and ∥ · ∥_1_ and ∥ · ∥_2_ denote the *L*_1_ and *L*_2_ norms respectively (Supplementary Information). For 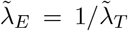, we showed that equation (4) becomes almost exactly a power law, namely 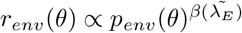. We derived an approximate analytic expression for the exponent, 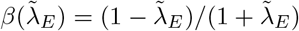, which matches the numerical results (Figure 4d; Supplementary Information). Equation (4) then becomes 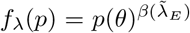. Consequently, by considering a pair of environments *A* and *B*, environment-invariance follows:

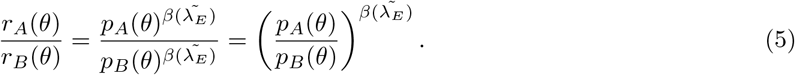

Environment invariance emerges as a consequence of the cost function considered to model adaptation in the power law regime.

### The model fits the power law in two environments and predicts the modulation of heterogeneous tuning curves in a new environment

The model predicted the adaptation of the experimentally recorded tuning curves, which are not homogeneous even in the uniform environment [1]. Inhomogeneities in tuning curves lead to a bias in the average population response to stimuli. The average population responses, therefore, are not power laws of the probability. In real data, thus, it is the ratio of average population responses to be a power law of the ratios of probabilities (Figure 1d). This is because taking the ratios of the average population responses cancels out the biases present in the empirical data. To verify whether the model is robust to these biases, we trained the model to fit the tuning curves adapted to two environments considered in [1], and tested if the model could predict responses to a third one. Specifically, through gradient descent, we trained the model to find a set of weights 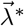 such that the adapted tuning curves were as close as possible to those of the data. For a specific recorded session, at gradient descent convergence (Suppl. FigureS.6a), we verified that the model was able to fit the data tuning curves (Suppl. FigureS.6b-c). Additionally, we verified the robustness of the fit by plotting the power law of the data and the power law of the fitted tuning curves of the model (Figure 5a), and observed a match between the two. Then we assessed if the trained model could predict the adapted tuning curves to the environment held out, by plotting the data tuning curves and the model’s prediction (Figure 5b), observing a good match. We repeated the procedure for all the recording sessions in [1]. The power law exponents in the model were compatible with those in the data (Figure 5c), and the *R*^2^ of the predicted tuning curves against the data (Figure 5d) was close to 1. The model is therefore able to predict the adaptation-driven changes in the single neurons’ heterogeneous tuning curves recorded in [1].

**Figure 5.**
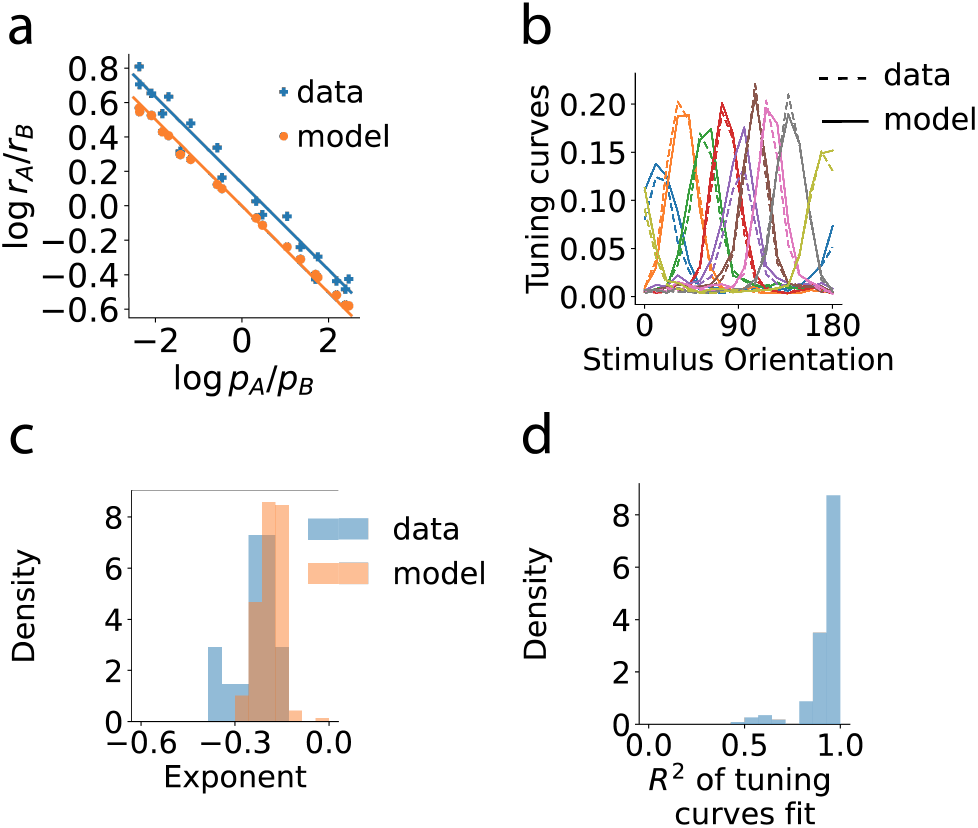
The model fits the heterogeneous tuning curves of the data, and predicts their changes due to adaptation. a,c) A fit of the data with the model. b,d) The predictions of the model on the data held out. A) Power law of adaptation in the data and corresponding fit from the model for a fixed recording session. b) At fixed recording session, tuning curves of the data adapted to a new environment (dashed) vs prediction of the model (solid). c) Exponent distribution for the data and for the model’s fit. d) Distribution of the *R*^2^ coefficient between the adapted tuning curves in the data and the model’s prediction.

### A competition between an orthogonality and a gain-difference terms for stimulus discrimination explains the change in power law exponent for natural stimuli

The exponent of the power law is invariant to the probability distribution of the stimuli shown, but it depends on the stimulus type [1]. The exponent is closer to zero for natural stimuli (−0.19 ± 0.03) than for oriented gratings stimuli (−0.25 ± 0.06). This raises the question of whether this difference could arise from differences in representation fidelity between the two stimulus types. We therefore first investigated whether neural representations differed between natural stimuli and oriented gratings, and then examined whether these differences could account for the observed change in the power-law exponent.

A key difference between neural responses to oriented gratings vs natural stimuli is a smaller minimal angle between neural responses to oriented gratings. We compared angles in the data of [1] between the high-dimensional neural representations (i.e., the response vectors) of natural stimuli vs oriented gratings (Figure 6a). While oriented gratings can be described by a single parameter (orientation angle), natural stimuli are inherently high-dimensional. As a result, a finite set of randomly sampled natural stimuli is likely to be highly dissimilar, whereas a finite set of oriented gratings will inevitably include some that are relatively similar to each other. This difference between stimulus types leads to differences in response vectors. Naming *x* a stimulus, we thus computed the angles between all pairs of response vectors *α*(*x*_*i*_, *x*_*j*_) to stimulus pairs *x*_*i*_, *x*_*j*_ from the data in [1]. Then, for each stimulus *x*_*i*_, we selected only pairs such that the other stimulus *x*_*j*_ formed the smallest angle in the neural space. Comparing the two distributions, we found that the minimal angle between response vectors, which we defined as *α*_min_, was higher if the stimuli were natural stimuli, rather than oriented gratings (Figure 6b; see Methods). This minimal angle is greater than zero due to the finite number of stimuli.

**Figure 6.**
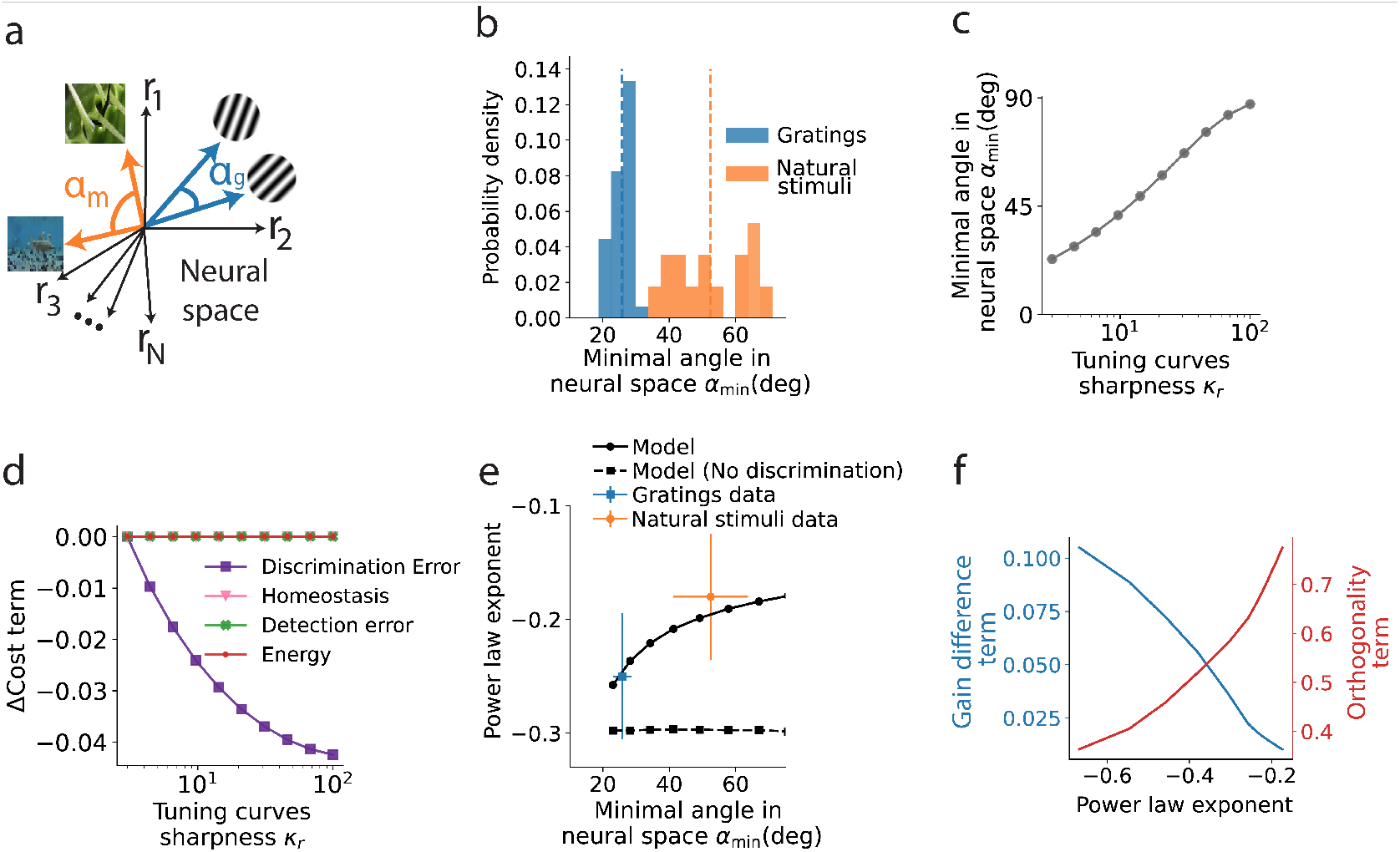
A reduction in discrimination error explains the change in power law exponent in natural stimuli. a) Outline of the angle between representations in the high-dimensional neural space identified as response vectors to a pair of oriented gratings (blue) and a pair of natural stimuli (orange). The subscripts *g* and *m* respectively stand for *gratings* and *natural stimuli*. b) Histogram of minimal angular distance between neural responses to a finite number of nearby oriented gratings (blue) and natural stimuli (orange), obtained by analyzing data from [1]. c) The minimal angle between neural response vectors to a fixed set of stimuli as a function of the tuning curves’ sharpness *κ*_*r*_ . d) Change, from a reference value at *κ*_*r*_ = 3, of the four cost terms as a function of the tuning curves’ sharpness *κ*_*r*_, for a fixed set of stimuli and a fixed probability distribution. e) Power law exponent as a function of the minimal angle between neural response vectors predicted by the model (solid black), in the gratings data (blue), in the natural stimuli data (orange) and as predicted by the model without the discrimination term (black, dashed). f) The exponent of the power law as a function of the two competing terms characterizing the discrimination error.

Increasing the minimal angle *α*_min_ between response vectors in the model decreased the discrimination error of the underlying reference tuning curves, leading to a monotonic decrease of the absolute value of the power law exponent. One way to model an increase of the minimal angle *α*_min_ between response vectors is to increase the sharpness *κ*_*r*_ of the tuning curves [53]. We thus changed the sharpness of the reference tuning curves and observed an increase of the minimal angle *α*_*min*_ (Figure 6c). While the discrimination error decreased, energy, detection error, and homeostasis constraint remained invariant with *κ*_*r*_ (Figure 6d). For fixed stimulus probability distributions, we optimized separately the cost function for distinct values of *κ*_*r*_ and found that increasing *α*_min_ from the value measured for gratings to the value measured for natural stimuli resulted in an increase in the power-law exponent compatible with that observed between the same two stimulus types(Figure 6e).

To show that the discrimination term is the one responsible for the change in the exponent, we excluded as a control the discrimination error from the cost function, and we saw no change in the power law exponent (Figure 6e). We identified a competition between two possible ways for achieving discriminability that explains the functional advantage of the change in power law exponent. The discrimination error can be decomposed into a gain-difference term and an orthogonality term (Supplementary Information): intuitively, at fixed *κ*_*r*_, discrimination can only be improved by increasing gain differences across stimuli, which is costly in energy, whereas increasing *κ*_*r*_ affords discriminability through reduced overlap between tuning curves. Therefore, the larger the sharpness *κ*_*r*_, the more the orthogonality term contributes to discrimination, allowing to save on the gain difference term (Figure 6f), resulting in a shallower power law. On the other hand, for lower *κ*_*r*_, discrimination can be achieved by having a strong modulation, and thus a steeper power law, to distinguish responses to different stimuli. Altogether, these results show that neural representations of natural stimuli, which are typically more mutually dissimilar than those of oriented gratings, reduce the gain modulation term for discriminability and thus reduce the steepness of the power law.

## Discussion

The discovery of the power law in [1], which links the probability of stimuli in an environment to the adapted average population response of sensory neurons, enables predictions about how neural responses adapt to new environments. In this context, we introduced an efficient coding model for adaptation that, for the first time, offers a normative explanation for the observed power law (Figure 1). We discovered that the environment-invariant power law reported in [1] can emerge from optimizing a tradeoff between energy of the neural code, its representation fidelity, and a homeostatic drive to baseline tuning curves (Figure 1). The experimentally observed invariance of the exponent to changes in environment naturally emerged from the efficient coding approach we developed, for a wide set of parameters (Figure 2), and we provided an analytic explanation for the emergence of the invariance (Figure 4). To explore whether an alternative trade-off between objectives could account for the experimental results of [1], we considered several combinations and definitions of coding objectives. Each of these combinations, however, was not able to match the main features of the data (Figure 3). Thus, to the best of our knowledge, our work provides a parsimonious, interpretable, testable model that explains the functional benefits of the power law and its related properties discovered in [1].

We validated the model against experimental data. Reproducing the universal power law served as an initial confirmation of the model. We also performed a new analysis of the data in [1] and discovered that the average modulation of the tuning curves during adaptation is primarily stimulus-dependent, a feature the model successfully captures. Additionally, by tuning the energy, detection, and discrimination weights in the cost function, we showed that the model could fit the heterogeneous tuning curves in experimental data. Using these tuned parameters, we then predicted the adapted response rates in a new environment (Figure 5).

We explained that the reduction in the magnitude of the power law exponent observed for natural stimuli (Figure 6) arises from the competition between two contributions to the discrimination error: a gain difference term, which dominates for oriented gratings, and an orthogonality term, which dominates for natural stimuli. We modeled the transition between oriented gratings to natural stimuli by increasing the sharpness *κ*_*r*_ of the tuning curves. Sharper tuning curves resulted in response vectors that were widely separated in angle, similar to those observed in natural stimuli data. As the angle between response vectors increased, we observed a smooth transition from the gratings exponent to the natural stimuli exponent.

The model is based on the efficient coding approach [24, 54, 55], namely that sensory neurons optimize the representation of the environment given a limited amount of available resources. Although it is widely accepted that neural processing is shaped by environmental statistics, quantitatively defining this relationship remains challenging [56]. Various theories have been proposed, such as sensory neurons maximizing mutual information between stimuli and their representations under limited coding capacity [36, 38, 51, 57], or aiming to eliminate redundancy in representations. Other perspectives suggest that the brain seeks to infer high-level hidden features of the environment within the constraints of a limited entropy rate [35], or that it minimizes the average reconstruction error while reducing metabolic costs [39]. The optimization process may also account for the role of downstream motor neurons [58] in the optimization process.

Our model is highly interpretable and we provided insight into the role of each term of the weighted objective function. The energy cost, detection error and homeostasis constraint are necessary and sufficient to get a negative power law exponent. The discrimination error is necessary to explain the presence of a neuron-dependent modulation of the tuning curves caused by adaptation and it explains the change in exponent for natural stimuli. By simplifying the weighted objective optimization, we derived an analytic solution for the optimal rate, which allowed us to interpret the emergence of the environment-invariance and the power law exponent.

Animals may dynamically adjust the relative magnitude of the weights, and thus the power law exponent, based on their energy-saving needs. This interpretation suggests testable predictions, namely that introducing a detection task for mice may shift the exponent toward a larger value. However, a change in behavior state does not necessarily change the exponent of the power law, as we showed by partitioning the data between rest and locomotion, which is known to amplify neural responses in the mouse cortex [48, 49].

Previous work by [1] explored the computational implications of the universal power law, characterizing how the strength of neural responses is related to stimulus surprise. It was noted that more surprising stimuli tend to require more energy to be represented. However, these findings are consequences that follow from the universal power law, while leaving open the question of why such a universal power law emerges and what adaptive role it serves. Our work addresses this question by introducing a weighted objective optimization problem whose solution gives rise to the universal power law.

Our model generates predictions that could guide future research. For example, future experiments may validate or refute the model’s prediction that an increased minimum angle between neural representations leads to a change in the exponent (Figure 6). One possible direction would be to investigate two-dimensional stimuli, such as gratings varying in both angle and spatial frequency, where we expect a greater minimal distance between a finite number of neural representations and a corresponding exponent closer to zero than in the one-dimensional case.

## Acknowledgments

This work was supported by NIH R21 EY035064 (DLR, MD, and MM). We thank Ramon Nogueira and Dean Buonomano for helpful comments on this manuscript.

## Author contributions

DLR, MD, and MM designed the model; MD and MM developed the model; MD and MM designed the data analysis; MM performed the data analysis; DLR, MD, MM, and SAM interpreted results; MD and MM wrote the original draft; DLR, MD, MM, and SAM edited the manuscript.

## Methods

### Sensory environments

Each stimulus was represented by an angle *θ* in the interval [0, *π*], corresponding to the orientation of a grating (in this context, an angle *θ* + *π* is equivalent to *θ*). A sensory environment *e* was defined by the probability density function (PDF) *p*_*e*_(*θ*) that the stimulus was presented. We studied continuous environments, namely the ones having a continuous PDF, and we discretized for computational reasons the domain [0, *π*] in *n*_*S*_ values in our numerical computations.

### Tuning curves

the model considered reference tuning curves *r*_0_(*φ, θ*) defined over the stimulus orientation *θ*, of neurons with preferred orientation *φ*. The reference tuning curves were modeled by von Mises functions:

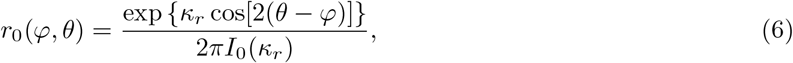

where *κ*_*r*_ measures the sharpness of the tuning curves and *I*_0_(*x*) is the modified Bessel function of the first kind of order 0. The factor 2 in front of the angular terms in the r.h.s. of the equation ensures that the function is periodic in [0, *π*] instead of [0, 2*π*].

### Weighted objective optimization

For a given environment *p*_*e*_(*θ*), the model finds the neuron-dependent and stimulus-dependent modulations, respectively *a*_*N*_ (*φ*) and *a*_*S*_(*θ*), such that the modulated tuning curves *r*(*φ, θ*) minimize the total cost in equation (2). The energy cost was defined as:

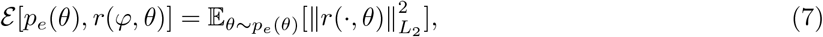

where:

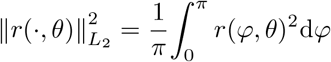

This quantity is related to the average population response over time, according to the idea that spiking consumes energy. The symbol E is the expected value. The subscript *θ* ∼ *p*_*e*_(*θ*) specifies that the average is with respect to the angular stimulus *θ*, which is stochastically drawn from the environment distribution of stimuli *p*_*e*_(*θ*):

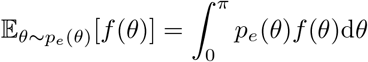

for a generic function *f* . The discrimination error was defined starting from the definition of d-prime [59–61]. As a simplifying assumption, we modeled the noise as independent, normal, and identically distributed across neurons, with a covariance matrix equal to I*σ*^2^, or, in an alternative formulation described in the Method section *Alternative model with Poisson statistics*, assumed the noise came from a Poisson distribution. For an independent, normal and identically distributed noise, the d-prime between the responses to two stimuli *θ*_1_ and *θ*_2_ is:

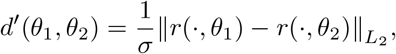

which can be interpreted as a signal-to-noise ratio, where the signal is the difference between mean responses. We defined the discrimination error by

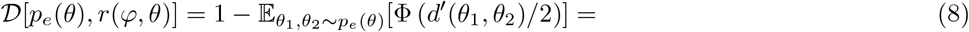

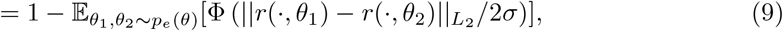

where the function Φ (*z*) is the cumulative distribution function for a normal distribution of zero mean and unit variance [59]:

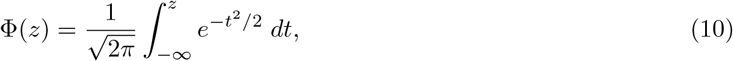

and where we averaged over pairs of stimuli, *θ*_1_ and *θ*_2_, drawn independently from the environment distribution *p*_*e*_(*θ*). The average over *θ*_1_ and *θ*_2_ should be interpreted as a time average over pairs of consecutive stimuli that the brain is trying to discriminate, and the cost defined thus represents the average discrimination error in the task.

The detection error was defined as:

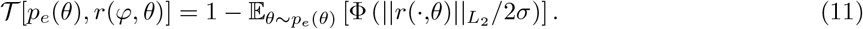

This term measures the error in the detection of a stimulus against a background. It is equivalent to the average error in the discrimination between the stimulus presented and another with zero contrast, which we model to elicit a response with zero amplitude.

The homeostasis constraint was defined as:

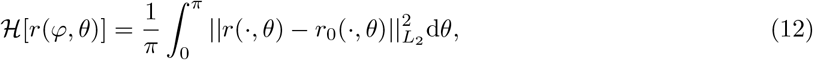

where *r*_0_(*φ, θ*) is the reference tuning curve defined in equation (6). This term penalizes adapted tuning curves that are too different from the reference ones [62]. Note that this form of homeostasis is different from the one described as a cause of sensory adaptation [50], which is instead defined as the invariance of the average firing rates across the population of neurons and across environments.

### The optimization process

We minimized the total cost *C* defined in eq. (2) using a gradient descent technique. We enforced a nonparametric optimization, discretizing the orientation space into *n*_*S*_ stimuli and the neural space into *N* neurons. The modulation functions, *a*_1_ and *a*_2_, were discretized as *a*_1_ = {*a*_1_(*φ*_1_), *a*_1_(*φ*_2_), …, *a*_1_(*φ*_*N*_ )} and 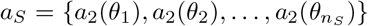. The number of optimized parameters was thus *n*_*S*_ + *N* . We initialized these terms to 1 and subsequently updated them via a gradient descent over the total cost. The gradient descent procedure was structured as follows:

- We computed *r*(*φ, θ*) = *r*_0_(*φ, θ*)*a*_1_(*φ*)*a*_2_(*θ*) for each *φ* ∈ {*φ*_1_, …, *φ*_*N*_ } and each 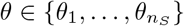
- We substituted the function *r*(*φ, θ*) onto the total cost *C* in equation (2) and we computed an auto-gradient numerically with respect to the *N* +*n*_*S*_ parameters. In the discretized form, each cost became:

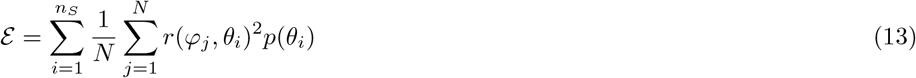

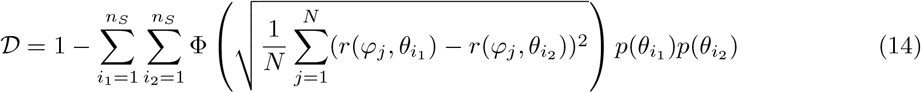

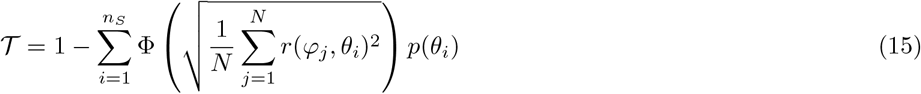

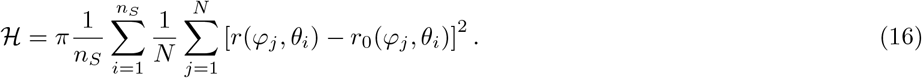

We have set the noise to a reference value, *σ* = 1*/*2. This is an arbitrary choice, which does not reduce generality, since the noise only gives a scale for the intensity of rates. By setting *σ* = 1*/*2, we are fixing such a scale.
- We updated each parameter by subtracting along the sign of the partial derivative, using a gradient descent method named Sign Descender [63], an effective method in numerical optimization problems [64].
- We repeated the procedure until convergence.

Once the optimal tuning curves were found, we computed the average population response as a function of the stimuli: 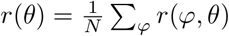. In our model we use average responses instead of their L2 norm, but the two are highly correlated in the data [1]. As a last step, we plotted log *r*(*θ*) as a function of log *p*(*θ*) and we linearly fitted log *r*(*θ*) as a function of log *p*(*θ*): log *r*(*θ*) = *β* log *p*(*θ*) + *q*. This linear relation implies the power law: *r*(*θ*) = *e*^*q*^*p*(*θ*)^*β*^ ∝ *p*(*θ*)^*β*^. Thus, by evaluating the score of the fit, it was possible to evaluate how close the relation was to a power law.

### Alternative models of adaptation

#### Covariance homeostasis model

We considered a model that enforces homeostasis at the level of pair-wise covariance between neuronal responses [2]. Specifically, we defined the covariance homeostasis cost as the *L*_1_ distance of the covariance from the reference covariance:

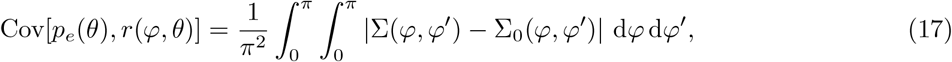

where the covariance matrices are given by:

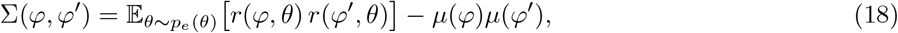

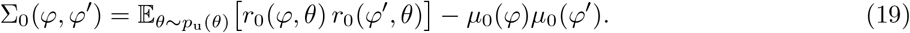

and the mean responses are defined as:

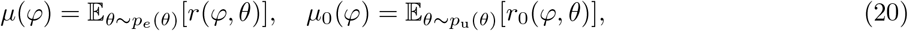

where *p*_u_(*θ*) denotes the uniform distribution over [0, *π*]. We additionally imposed the homeostasis constraint on tuning curves defined in equation (12), as in [27]. The total cost function was thus defined as:

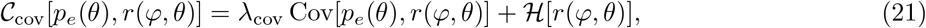

where *λ*_cov_ is a weighting parameter and ℋ is the homeostatic term defined in equation (12).

#### Product homeostasis model

As a related alternative, we considered a model that enforces homeostasis on second-order response moments without subtracting the mean activity [2]. We defined the product homeostasis cost as the *L*_1_ distance of the response product from the reference product:

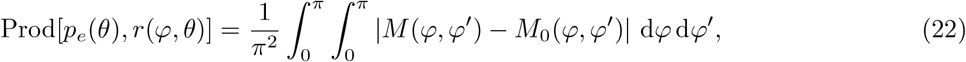

where the second-order moment matrices were defined as:

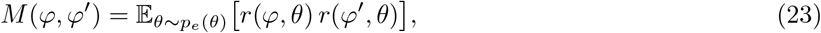

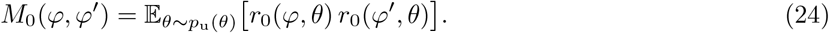

The total cost function was defined as:

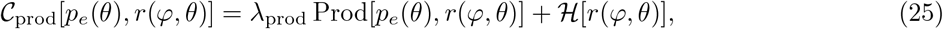

where *λ*_prod_ is a weighting parameter.

#### Mutual information maximization

We considered a model that maximizes an approximation of the mutual information between stimuli and neural responses. The cost function is defined as:

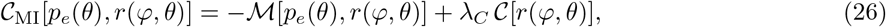

where *λ*_*C*_ is a weighting parameter, ℳ is a proxy for the mutual information expressed in terms of the Fisher information,

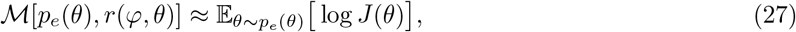

where *J*(*θ*) is the Fisher information, and *C* is a constraint on the total Fisher information capacity [38]:

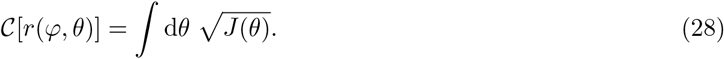

Assuming independent Gaussian noise with variance *σ*^2^, the Fisher information is given by [47]:

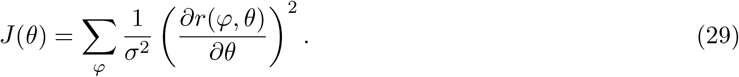

The derivative with respect to *θ* was computed numerically using a central finite-difference scheme with periodic boundary conditions, reflecting the circular structure of the stimulus space.

#### Reconstruction error minimization

We considered a model that minimizes a reconstruction error functional defined as [51]:

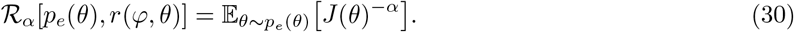

where *α >* 0 is a parameter controlling the strength of the penalty on low Fisher information values. In our simulations, *α* was fixed to 1, corresponding to the discrimax case [37]. The cost function was defined as

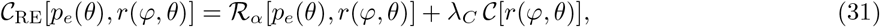

where *λ*_*C*_ is a weighting parameter (equation (28)).

#### Information maximization under constraints on energy

We considered a model that maximizes a power of the Fisher information under the constraints of limited energy and Poisson noise statistics [37]:

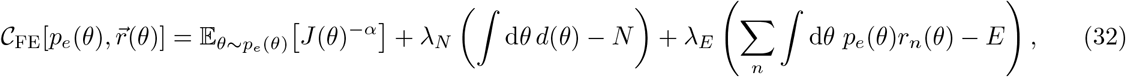

where the quantity *d*(*θ*) is the density of neurons with preferred orientation *θ*. The second addend ensures that the total number of neurons remains constant. The term is needed because the model allows changes in the density of neurons’ preferred orientation and on a neural modulation *g*(*θ*). Specifically, the tuning curve of a neuron *n* was parameterized as:

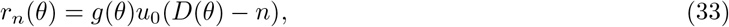

where *D*(*θ*) is the cumulative of the density and *u*_0_ is a von Mises function with peak at 0:

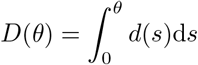

with *D*(*π*) = *N* . The optimal density and gain are (Supplementary information and [37])

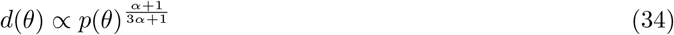

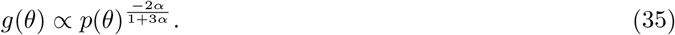

We plugged these functions in equation (33) to extract the adapted tuning curves values. Then we numerically computed the average firing rate as a function of the probability and fitted a power law. To achieve an exponent of -0.25, we set *α* = 0.18.

#### Best hyperparameter selection

We selected, for each model, the parameter configuration that best matched the target phenomenology, namely high *R*^2^, high *ω*, and an exponent close to *β*_data_ = −0.25. This was achieved by minimizing the following objective:

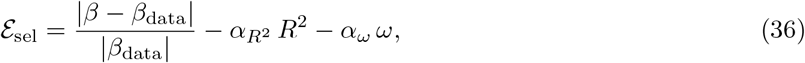

where 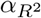 and *α*_*ω*_ are positive weighting constants. In practice, we assigned a stronger weight to the *R* term in order to prioritize solutions that closely followed a power-law relationship. In particular, 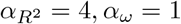.

### Assessment criteria

For every model we considered, we performed a grid search over the corresponding hyperparameters. Unless otherwise specified, the grid spanned values logarithmically spaced between 10^−3^ and 10^3^. For the covariance homeostasis model, we explored the parameter *λ*_cov_ and the reference baseline for the covariance. Similarly, for the product homeostasis model, we explored the parameter *λ*_prod_ and the reference baseline for the responses product. For the mutual information model and for the reconstruction error model, we explored the parameter *λ*_*C*_. For each combination of parameters, the tuning curves were optimized using the same procedure adopted for the full model described above. So we looked for the modulations *a*_1_(*φ*) and *a*_2_(*θ*) such that the modulated tuning curves *r*(*φ, θ*) = *a*_1_(*φ*)*a*_2_(*θ*)*r*_0_(*φ, θ*) optimize the cost function defined for each model. To compare models, we evaluated three quantities: (i) the coefficient of determination *R*^2^ of the fit, (ii) an invariance score *ω*, and (iii) the exponent *β* of the fitted power law.

#### Assessment of fit

We used the coefficient of determination (*R*^2^) throughout the paper to quantify the quality of each fit. Specifically, we evaluated *R*^2^ for the linear fit of log[*p*_*A*_(*θ*)*/p*_*B*_(*θ*)] versus log[*r*_*A*_(*θ*)*/r*_*B*_(*θ*)], as well as for the model fits of the neural tuning curves *r*(*φ, θ*) (Figure 5b).

#### Invariance measure

The invariance *ω* quantifies how consistently response curves obtained under different environments collapse onto a single functional relationship when expressed as a function of the stimulus probability. For each pair of environments *e*_1_ and *e*_2_, we considered the curve 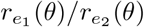 (firing rates) as a function of the stimulus probability ratio 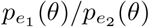. These curves are first restricted to their common domain [*p*_ratio min_, *p*_ratio max_], defined as the intersection of the supports across environments pairs. Each curve is then interpolated over a shared grid 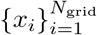 in this interval. Denoting the interpolated curves as 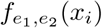, we define the mean curve

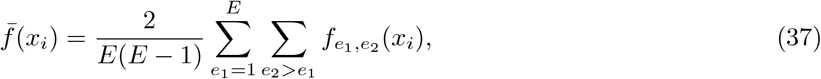

where *E* is the number of environments. The invariance is then defined as the average coefficient of determination of each curve with respect to the mean curve:

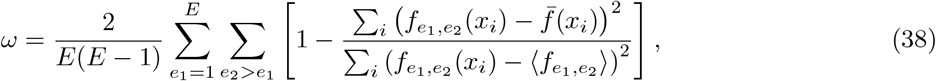

where 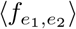 denotes the average of 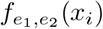 over *i*. This quantity takes values close to 1 when all curves collapse onto a common function, and decreases as variability across environments increases.

### Poisson statistics

Given two stimuli, *θ*_1_ and *θ*_2_, each presented for a duration *T*, each neuron produces a spike train that depends on both the stimulus and the environment, and that is subject to stochastic variability due to noise. We model each neuron’s spike train as a homogeneous Poisson process with a stimulus-dependent rate. Let *n*_*i*_(*θ*) denote the total number of spikes emitted by neuron *i* during the presentation window *T* of stimulus *θ*. Then *n*_*i*_(*θ*) is Poisson distributed with mean *r*_*i*_(*θ*)*T* . We can define the empirical firing rate as:

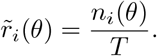

This is a random variable with expected value equal to the expected rate *r*_*i*_(*θ*), and variance equal to 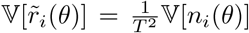. Due to the fact that *n* _i_ (*θ*) is Poisson distributed, then 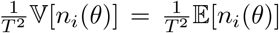, so 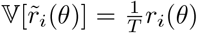. It is possible to show that using the central limit theorem, the discrimination cost is (Supplementary information):

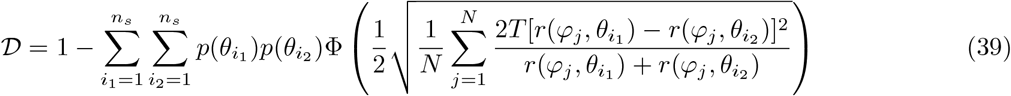

where *T* was fixed such that the reference tuning curves had identity matrix covariance (Supplementary Information). Similarly, the detection cost is (Supplementary information):

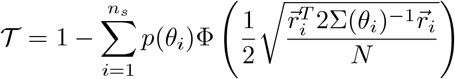

where:

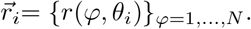

### Data analysis

We analyzed data from [1], which consisted of 20 recording sessions from four adult mice in the mouse primary visual cortex (V1). In each recording session, a head-fixed mouse passively viewed oriented gratings while running on a treadmill. Neural responses from hundreds of V1 neurons were simultaneously recorded using calcium imaging. The stimuli were sets of 18 oriented gratings or 18 natural stimuli, each displayed on a screen for 300 ms. Each recording session included six blocks of approximately 100 stimuli. Within a block, stimuli were drawn from three smaller blocks randomly permuted, each one corresponding to a different environment.

#### Analysis of modulation in experimental data

To determine the structure of the adaptation-induced modulation (Figure 1e), we considered only recording sessions where one of the environments was uniform. We then used these recordings to compute the non-adapted average tuning curves of neurons with similar preferred orientations. We reduced the original set of neurons to a set of *N* = 18 super-neurons [39, 50, 65]. These super neurons are defined by taking the average of all neurons whose preferred orientation was within a given range of orientations. The orientations defining the borders of these ranges were evenly spaced within [0, *π*]. We computed the modulation by taking the ratio of the super-neurons tuning curves, *a*(*φ, θ*) = *t*_*e*_(*φ, θ*)*/t*_*U*_ (*φ, θ*), where *U* stands for the uniform environment, *e* ∈ {*A, B*} is one of the two biased environments and *t*_*x*_(*φ, θ*) are the recorded tuning curves for environment *x*. All the biased environments of these selected recording sessions followed a von Mises distribution with the same sharpness, differing only in the position of the peak. This symmetry allowed us to average the modulation across biased environments and recording sessions by applying a circular shift to the modulation to align each one with the same reference stimulus and preferred orientation.

#### Analysis of the power law under different locomotion states

We partitioned the data from [1] based on the mouse’s speed. Since mice were free to run on a treadmill, their speed varied continuously; we set a threshold of *v*_th_ = 1 cm*/*s to classify periods as rest (*v < v*_th_) or running (*v* ≥ *v*_th_). For each recording session and environment *e*, we constructed two response vectors per stimulus 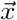, corresponding to the two locomotion states: 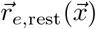 and 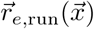. We then treated these vectors as independent and compared both their magnitudes and the exponent of the corresponding power law. A challenge arises because some sessions were dominated by one locomotion state, leading to too few observations for certain stimuli in the under-represented state. In such cases, these stimuli were discarded when their probability of arising was less than 1%, and the power law was fitted using the remaining data.

### Fitting of the data tuning curves and prediction of responses to a new environment

Each recording session in [1] has responses to three different environments. We considered, for a fixed recording session, two out of the three environments and we found the best weights 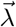 to reproduce the recorded neural responses. Specifically, we selected a training environment pair A and B. Denoting by *t*(*φ, θ*) the recorded tuning curves from [1], averaged by neurons belonging to the same bin of 10 degrees of preferred orientation, we built the reference tuning curve as an average of the responses normalized across stimuli:

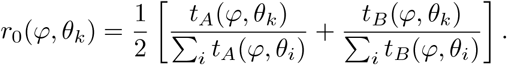

We initialized the weight vector to 10 different random initial conditions to study if the results converged in a robust way. At fixed 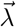, we minimized the total cost in equation (2) and obtained the adapted rates predicted by the model, 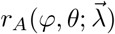 and 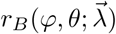, where the dependence on the weight vector 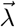 is explicitly shown. We then updated the weight vector 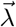 by descending the gradient of the error between the model’s tuning curves and the recorded tuning curves:

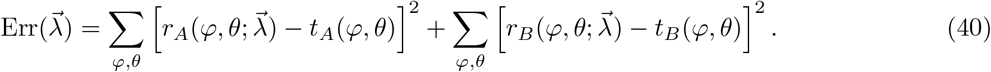

Specifically, we evaluated the gradient numerically, via finite difference, and used it to update 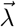 using the Sign Descender method [63]. We used the updated weight vector to compute the tuning curves optimizing the cost. We stopped the procedure when both the weight vector and the error in the tuning curves converged. The optimal 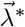, the one minimizing the error, was then used to predict the responses in the third environment, which we compared to the recorded responses from the experiments in [1].

### Angular spacing between responses

Let us consider the response vector **r**(*x*) = {*r*_1_(*x*), *r*_2_(*x*), …, *r*_*N*_ (*x*)} to a stimulus *x*, which could represent either an oriented grating or a natural stimulus. Each element of the response vector corresponds to a different neuron rate among the *N* neurons recorded. We computed the angle between the response vectors to two different stimuli *x*_1_ and *x*_2_ as:

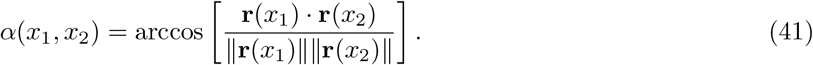

We used such a computation in the study of the difference in exponent between gratings and natural stimuli (Figure 6). In particular, we characterized the distribution of minimal angles between response vectors to natural stimuli and oriented gratings. To do so, we considered, for each stimulus *x*_1_, the stimulus *x*_2_ such that the angle *α*(*x*_1_, *x*_2_) was minimal. We iterated through the five recording sessions of natural stimuli in [1], and for each of them we iterated through the three environments recorded. We built the histogram of all *α*_min_(*x*_1_, *x*_2_) across recording sessions and environments in Figure 6b.

### Change in tuning curve sharpness

To model responses to natural stimuli, we increased the minimal angle between response vectors by increasing the sharpness *κ*_*r*_ of the von Mises reference tuning curves *r*_0_(*φ, θ*) and multiplying by a pre-factor *C*_0_(*κ*_*r*_) such that that the *L*_2_ norm of *r*_0_ is maintained constant as a function of *κ*_*r*_:

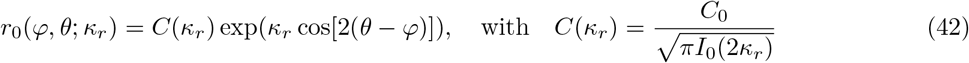

with such a change, we were able to model the change of exponent due to a change in orthogonality and gain-difference terms (Supplementary information).

## Supplementary information

### The reconstruction error model gives positive power law

The optimal fisher information is

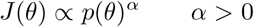

as shown in [51]. The Fisher information *J*(*θ*) can be expressed as function of the tuning curves *r*(*φ, θ*) = *a*(*φ, θ*)*r*_0_(*φ, θ*). For the case of gaussian isotropic noise [47]:

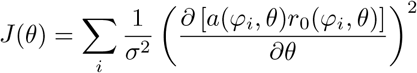

Imposing (close to what the data shows [1]) *a*(*θ, φ*) = *a*(*θ*), so stimulus modulation only, the Fisher information is:

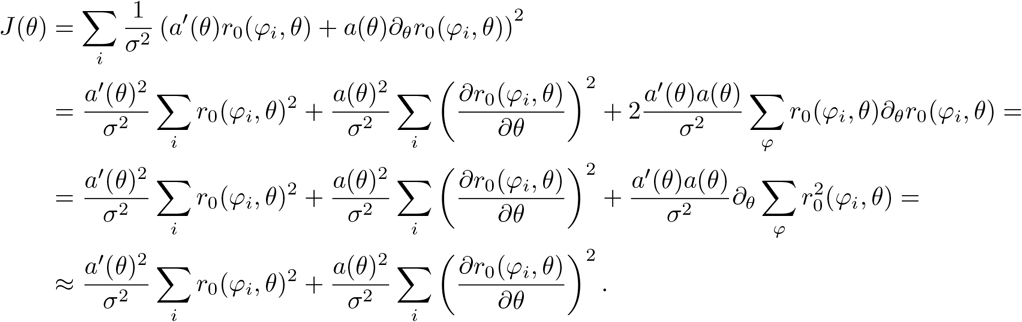

where we have assumed a dense tiling of neurons [37] to make the third addend vanish. By identifying the terms in the sums we obtain:

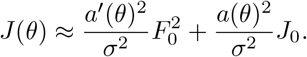

The optimal Fisher information is a power law [51]:

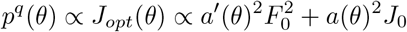

To get an invariant power law the term 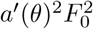 must be negligible compared to *a*(*θ*)^2^*J*_0_, otherwise a paradox emerges: suppose indeed that *a*(*θ*) ∝ *p*(*θ*)^*β*^. Then we see:

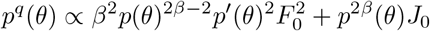

This equation cannot be solved for an arbitrary *p*. To get invariance, *a*(*θ*) and *r*_0_(*φ, θ*) must be such that:

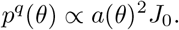

When this is the case, then the power law exponent is:

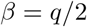

Since *q >* 0 in order to ensure maximization of the Fisher information, then the exponent of the power law must be positive, which is in contradiction with the negative exponent *β* = −0.25 observed in the data of [1].

## Cost function and its optimum for the model of information maximization under constraints on energy

To intuitively see why *d*(*θ*) is a density, let’s observe that a neuron’s preferred orientation is *θ*_*n*_ such that *D*(*θ*_*n*_) = *n*, because this vanishes the argument of the von Mises function *u*_0_. The preferred orientation of neuron *n* is thus *D*^−1^(*n*). So, in an interval in which *d*(*θ*) is large, the cumulative *D*(*θ*) explores a wide range of values, implying that its inverse *D*^−1^ maps into that interval many values of *n*. With the parameterization described above, it is possible to rewrite the cost function [37]:

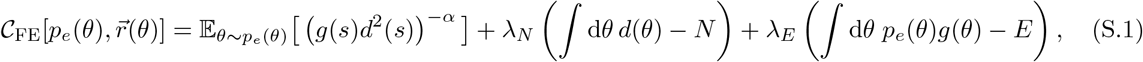

which are minimized analytically in [37], getting:

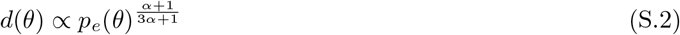

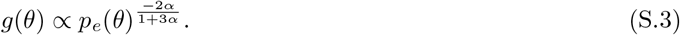

### Discrimination and detection costs under Poisson statistics

Using the Central Limit Theorem, the distribution of 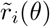 can be approximated by a Gaussian with moments determined in the main text [66]:

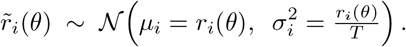

Therefore, the response vector 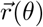 has a neuron- and stimulus-dependent noise. For a given stimulus *θ*, the covariance is in the form of a diagonal matrix:

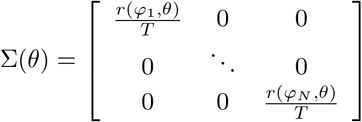

Given two stimuli *θ*_1_ and *θ*_2_, the exact decision boundary between the responses elicited by the two stimuli is quadratic, because the covariance is non-constant across stimuli [67]. We considered the approximation based on the average of the two covariance matrices [67]:

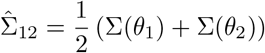

so that the discrimination accuracy is given by the Linear Discrimination Analysis formula [59]:

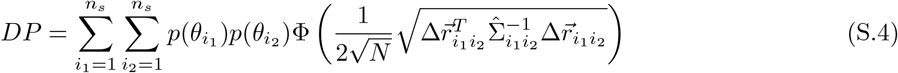

where 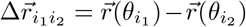. This linear decision boundary approximation could plausibly be implemented by downstream neurons acting as a linear decoder, with weights corresponding to their pre-synaptic projections. The matrix 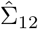 is also diagonal, because it is the sum of two diagonal matrices. Thus, its inverse is diagonal too:

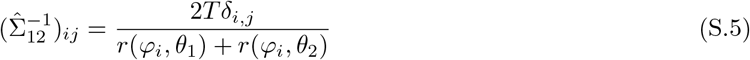

where *δ*_*i,j*_ = 1 if *i* = *j* and *δ*_*i,j*_ = 0 if *i j*. The DP is thus:

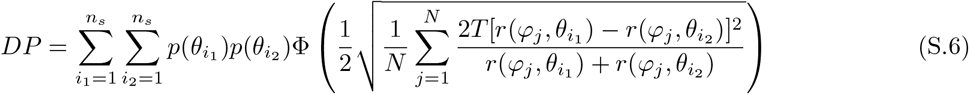

where we fixed *T* as the mean of the tuning curves over neurons and stimuli, i.e. such that for the reference tuning curves and the uniform environment the average covariance matrix is the identity: 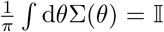. The detection cost follows the same logic and approximations described for the discrimination error, with the key difference that it measures the discriminability between a stimulus and the background.

### Analytic approach for a simplified model

We considered a simpler optimization problem than the one considered in equation (2) and we solved it analytically, showing that a power law emerges across a broad region of parameter space. This simplified model optimizes only energy cost, detection error, and homeostasis constraint:

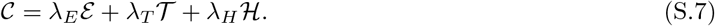

We are including the weight *λ*_*H*_ in this model, without setting it to 1 as in the main text, to study the limit of *λ*_*H*_ *>> λ*_*E*_, *λ*_*T*_ . To analytically tackle the optimization of this function, we first introduced two simplifying hypotheses:

1. In the detection error, defined in equation(11), we substituted Φ(*z*) with the identity *z*, so that the detection error became:

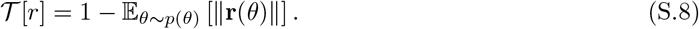

Since the noise is set to the reference value, this formula corresponds to one minus the average signal-to-noise ratio of responses.
2. We imposed that the modulation is only dependent on the stimulus:

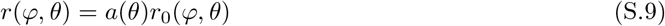

where *r*_0_(*φ, θ*) are the reference tuning curves, and *a*(*θ*) is their modulation due to adaptation. The dependence of the modulation on the stimulus alone is an approximation, justified by the fact that the modulation in the full model is mainly dependent on the stimulus (Figure 1e-j).

We considered a discretized form of the total cost to compute its optimization by means of partial derivatives rather than the functional derivative. We discretized the continuous stimulus space into *n*_*S*_ values, 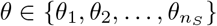. The probability of stimuli thus becomes 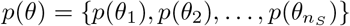, and the modulation becomes 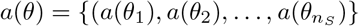. Therefore, the number of stimuli *n*_*S*_ represents the total number of discretization steps. The total cost for the simplified model is thus:

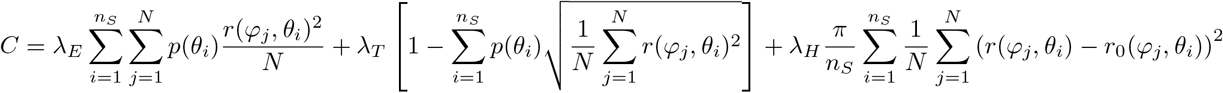

where we have also discretized the number of neurons into *N* values *φ* ∈ {*φ*_1_, *φ*_2_, …, *φ*_*N*_ }. Using equation (S.9), the total cost becomes:

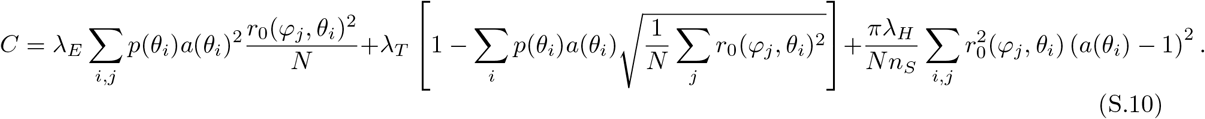

We define now:

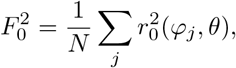

which is independent of *θ* for the reference tuning curves when the number of neurons *N* is large:

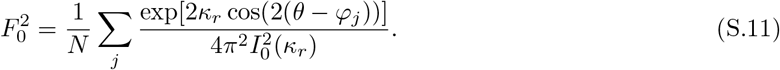

Defining d*φ* = *π/N* :

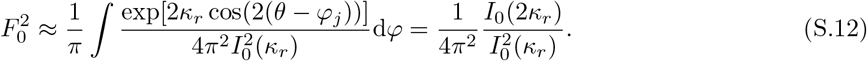

Inserting *F*_0_ in the total cost, eq. (S.10), we obtain:

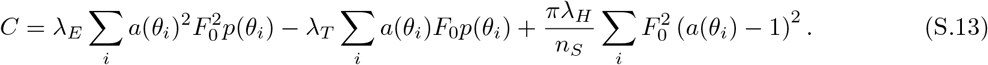

To find the local minima of the total cost of eq. (S.13), we set to zero the partial derivative with respect to a generic element *a*(*θ*), with 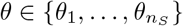:

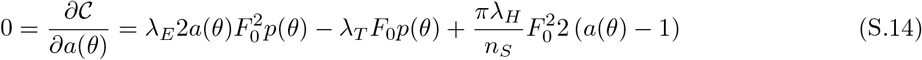

And thus we have:

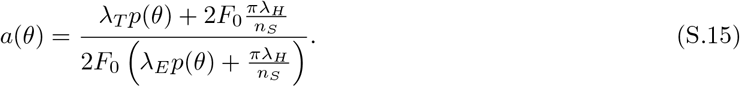

This is the expression for the modulation. Notice that if the homeostasis weight dominates the others, then the modulation tends to 1. This implies that the adapted tuning curve is identical to the reference tuning curve. If no homeostasis is instead present (*λ*_*H*_ = 0), the modulation is a constant whose value is fixed by a trade-off between energy and detection weights. Notice also that the probability of a stimulus scales with the inverse of the total number of discretization steps *n*_*S*_. Defining a probability density *p*(*θ*) = *f* (*θ*)*/n*_*S*_, the modulation loses its dependence on *n*_*S*_:

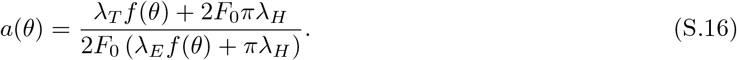

In a regime in which *λ*_*E*_ ≫ *λ*_*H*_ ≫ *λ*_*T*_, the law is a power with exponent −1. We can rewrite the modulation in a more compact form:

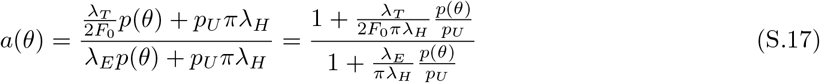

in which we have factored out 2*F*_0_ and substituted *p*_*U*_ = 1*/n*_*S*_. The average population response is proportional to the modulation, under the hypothesis of stimulus-dependent modulation of equation (S.9):

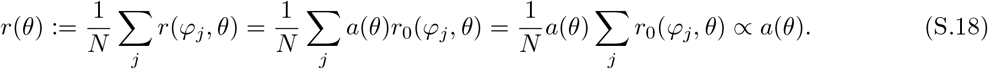

Equation (S.18) implies that if the modulation is a power of the stimulus probability, then the rate is a power of the probability too, *p*(*θ*)^*β*^ ∝ *a*(*θ*) ∝ *r*(*θ*). In the next section, we show that the modulation is invariant under environment changes and approximately equal to a power law for an extended region of the parameters.

### The relation between average population responses and probabilities is invariant under environment change

Consider equation (S.17). The modulation *a*(*θ*) does not directly depend on the stimulus value, but only on its probability. We can thus rewrite the modulation as a function of the probability of stimuli:

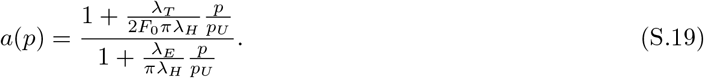

The functional form of *p*(*θ*), thus, is irrelevant to determine the modulation. This implies that the modulation is invariant to environment probability changes. The average population response is also invariant to environment probability changes, since it is proportional to the modulation, see eq. (S.18). This proportionality holds only under the simplifying assumption that the modulation is mainly stimulus dependent.

### Approximate power law for an extended region of the parameter space

In the reduced model (energy-detection-homeostasis, eq (S.10)), the optimal modulation approximates a power law across a range of stimulus probabilities consistent with experiments and over an extended region of the weight space. To demonstrate this, we studied for which regime equation (S.19) approximates a power law. We reformulated equation (S.19) by defining 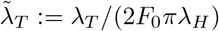 and 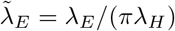:

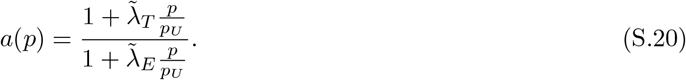

Then, we studied the values of 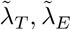 for which equation (S.20) is approximately a power law. We thus looked for the power law *cp*^*β*^ best approximating *a*(*p*), by minimizing the following sum of squares error:

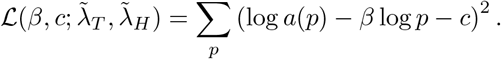

We numerically evaluated the error ℒ^∗^ for the optimal power law approximating the modulation law in equation (S.20) in the space 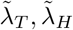. We found from the numerical analyses that the error ℒ^∗^ for the optimal power law is low along a line constrained by log 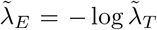 (Figure 4a-b), namely for 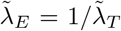. To gain further insight into this result, we rewrote the logarithm of the modulation log *a*(*p*) as a function of the logarithm of the probability ratio log 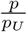, and we noticed that under the constraint 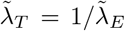, the quantity log *a* is an odd function of log(*p/p*_*U*_ ) up to a constant, consequently vanishing the quadratic term in its Taylor expansion. In fact:

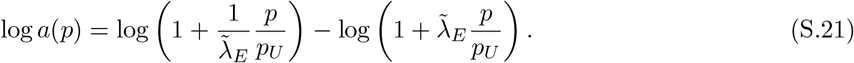

Defining 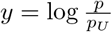 and rewriting the logarithm of the modulation as a function of *y*, i.e. *f* (*y*) = log *a*(*p*):

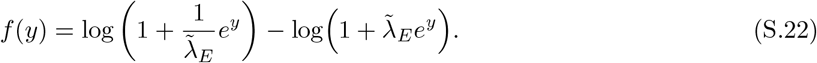

Now we can observe that 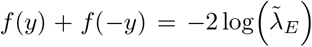, so that 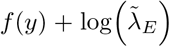 is odd, so that the even derivatives (except for the order zero) vanish . Therefore, expanding the modulation around *y* = 0 (i.e., *p* = *p*_*U*_ ) we have:

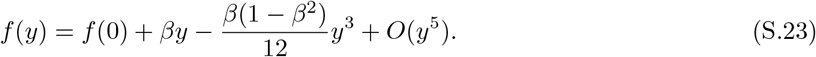

where 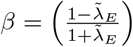 and we have dropped for sake of compactness the dependence of *β* on 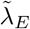. The expansion variable 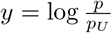 quantifies the degree with which the probability distribution *p* deviates from the uniform distribution. If we assume that such a deviation is small, we can truncate the Taylor series at its first order on *y*. Substituting 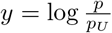 and *f* (*y*) = log *a*(*p*), we obtain:

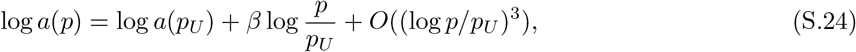

Evaluating the exponential of both sides of the equation we get:

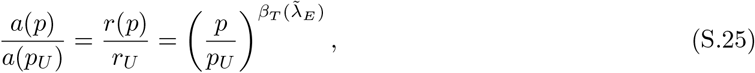

where we have used that *a*(*θ*) ∝ *r*(*θ*) and where:

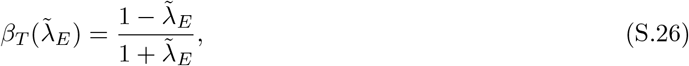

with the subscript *T* indicating that the result follows from the Taylor expansion and where we reintroduced in the notation the explicit dependence on 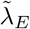. We compared the result for the exponent of the Taylor expansion with the exponent of the best power law fitting the modulation and we found almost perfect agreement (Figure 4d).

### The power law exponent does not change with sharpness when removing the discrimination term

We numerically observed that, upon removing the discrimination error, the power-law exponent remains independent of *κ*_*r*_. This can be understood analytically by noting that the remaining cost terms, energy, detection error, and homeostasis, are approximately independent of *κ*_*r*_. As a consequence, a solution that minimizes the cost for a given *κ*_*r*_ remains optimal for all values of *κ*_*r*_. To show this, we begin with a useful approximation. Consider tuning curves of the form:

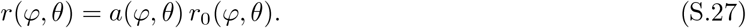

The *L*_2_ norm of the response at fixed *θ* is:

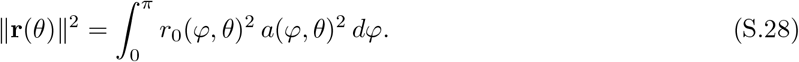

In the regime in which the reference tuning curves vary more rapidly than the modulation *a*(*φ, θ*), since *r*_0_(*φ, θ*) is sharply peaked around *φ* = *θ* with width *ε*, we approximate:

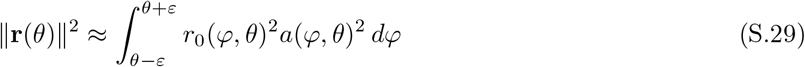

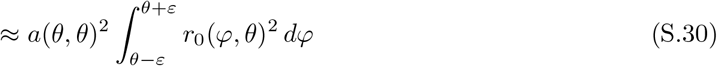

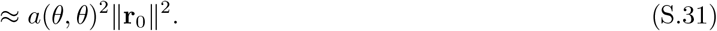

Thus, the norm of the response is effectively independent of *κ*_*r*_. We validated this approximation numerically by optimizing the full cost for every *κ*_*r*_ and compared ∥**r**(*θ*)∥^2^ with its approximate version *a*(*θ, θ*)^2^∥**r**_0_∥^2^. As expected, the approximation improves with *κ*_*r*_, but it’s already very good at *κ*_*r*_ = 3, with an error of only 2.5% (Suppl. Figure S.7). We now analyze the cost terms. The energy is:

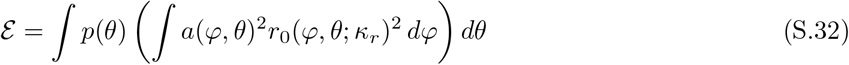

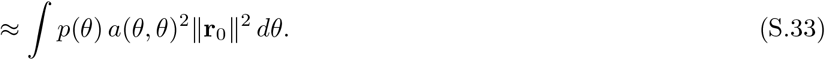

which is a constant w.r.t. *κ*_*r*_. The detection error is:

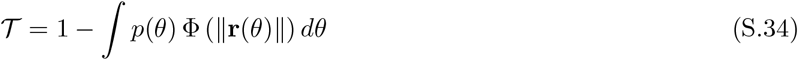

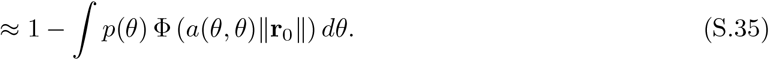

The homeostasis constraint is:

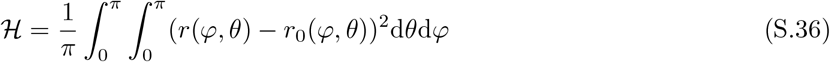

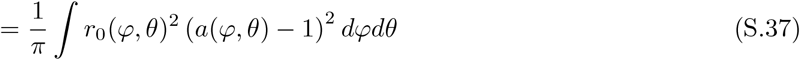

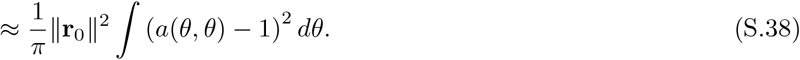

All these terms are approximately independent of *κ*_*r*_, proving that the power law exponent is constant in *κ*_*r*_.

### Effect of the discrimination term on tuning curves

The discrimination term is

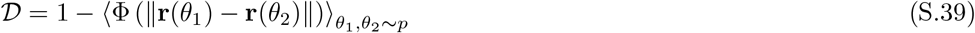

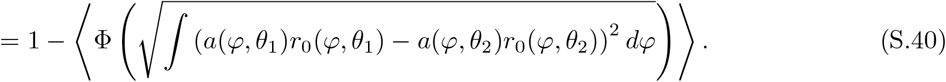

Expanding the square,

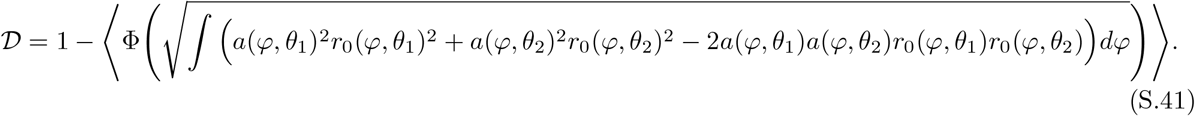

Using the previous approximation,

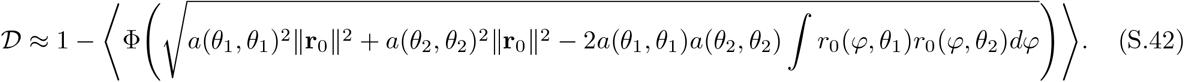

For fixed *θ*_1_ and *θ*_2_, the first two terms are independent of *κ*_*r*_ because ∥*r*_0_∥^2^ is independent of *κ*_*r*_ by construction. The cross-term, instead, decreases as *κ*_*r*_ increases, since it is a scalar product between tuning curves becoming increasingly orthogonal with growing *κ*_*r*_. As a result, the discrimination error decreases with increasing *κ*_*r*_. Therefore, unlike the other cost terms, the discrimination term introduces an explicit dependence on *κ*_*r*_, which in turn affects the optimal tuning curves and the resulting exponent. The pairwise squared distance entering *D* can be decomposed exactly (by completing the square) as

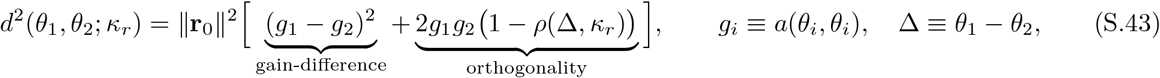

where *ρ*(Δ) = ⟨**r**_0_(·, *θ*_1_), **r**_0_(·, *θ*_2_)⟩*/*∥**r**_0_∥^2^ is the normalized overlap between reference tuning curves at two different stimuli, computed directly from the discretized, normalized tuning curves used in the model. Discriminability between *θ*_1_ and *θ*_2_ can thus be achieved through two mechanisms: by making the local gains *g*_1_, *g*_2_ different from one another, or through orthogonalization of the tuning curves themselves (*ρ* → 0), which is entirely determined by *κ*_*r*_. At fixed small *κ*_*r*_, tuning curves broadly overlap, so discriminability can only be achieved through the gain-difference term; because ℰ penalizes gain most strongly where *p*(*θ*) is large, the cost-minimizing solution suppresses *g*(*θ*) at the peak of *p*(*θ*) and allocates gain to the tails, producing a power law *g*(*θ*) ∼ *p*(*θ*)^*β*^ with *β* strongly negative. As *κ*_*r*_ increases, *ρ*(Δ) → 0, and the orthogonality term guarantees discriminability. The gain-difference term is then no longer needed to satisfy *D*, and the term reaches its maximum, so 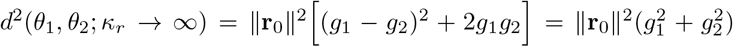, which optimizer relaxes toward a flatter solution favored by ℰ and ℋ. For the case of l*κ*_*r*_ → ∞, the orthogonality under the approximation of slow varying modulation gives *d*^2^(*θ*_1_, *θ*_2_; *κ*_*r*_ → ∞) ≈ ∥**r**_1_∥^2^ + ∥**r**_2_∥^2^. So when the orthogonality term saturates, discrimination can still be improved by increasing the overall responses to stimuli. In this regime, discrimination works in the opposite way of the energy term.

---

**Figure S.1.**
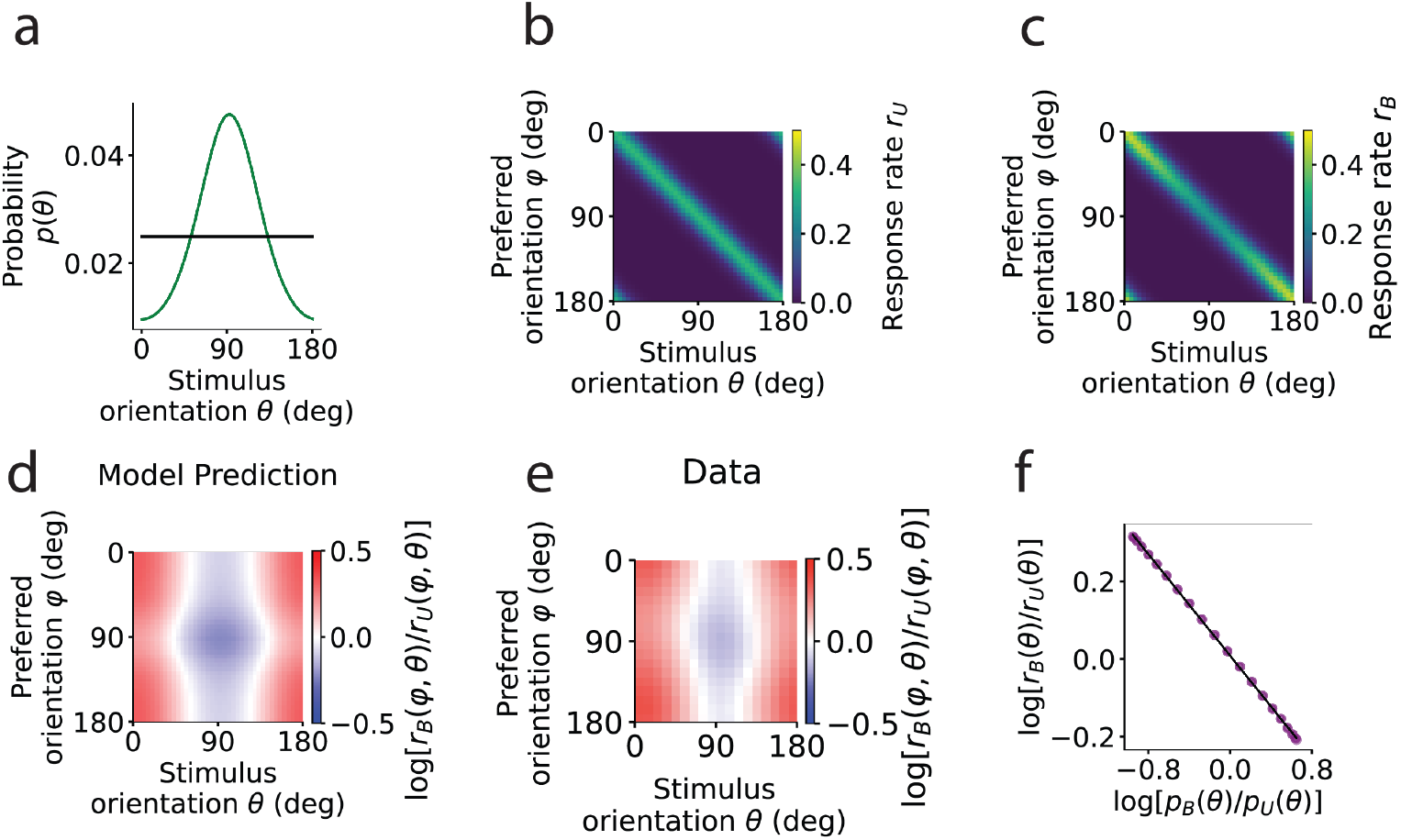
The model can reproduce the power law under Poisson noise statistics. a) Two example environments: a uniformly distributed environment (black) and a von Mises distributed environment (green). b) Tuning curves, represented as a heat map depicting the response rate as a function of stimulus *θ* and neuron’s preferred orientation *φ*, emerging from optimization of the total cost in equation (2), in the uniform environment in (a). c) Similar to (b) but the environment is the von Mises in panel (a). d) Modulation of the tuning curves emerging from the model as the ratio between the tuning curves in (c) and in (b). e) Modulation in experimental data in [1] obtained by averaging groups of neurons with similar orientation preferences. f) Logarithm of the ratio of average population responses between the von Mises and uniform environments in (a) as a function of the logarithm of the ratio of their probabilities (circles) and the power law fit (solid line).

**Figure S.2.**
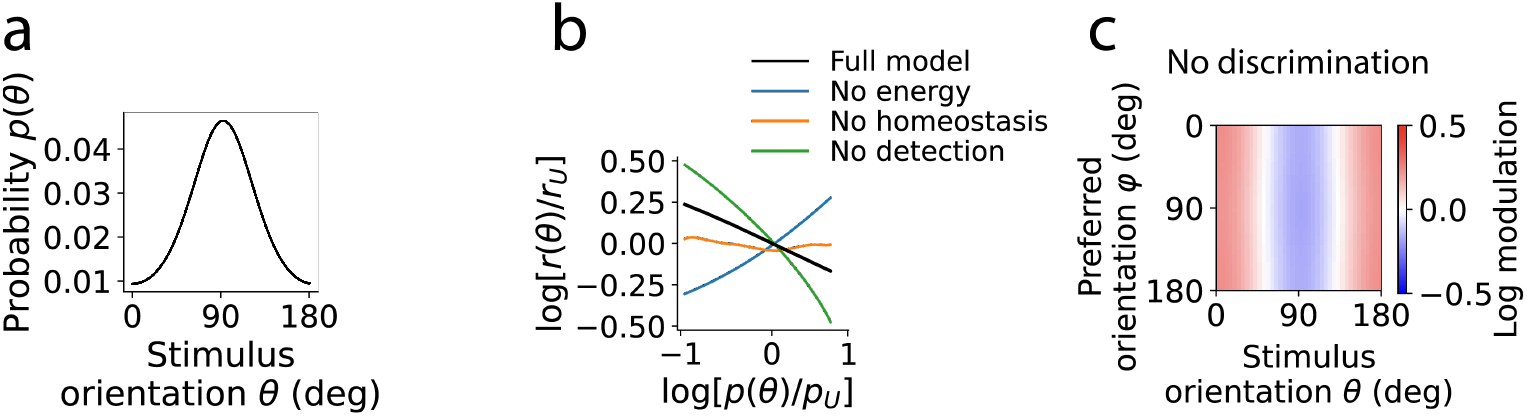
Energy, discrimination, detection and homeostasis are all required to account for the key experimental findings. a) Probability distribution of stimulus orientation defining a sensory environment. b) Adaptation to environment in panel (a) predicted by the full model or by a modified model with one weight set to zero at a time. Logarithm of the ratio between the response rates in the biased and uniform environments, against the logarithm of the ratio between the corresponding stimulus probabilities. c) Same as Figure 1j, but for the model in which discrimination error has been removed. Compare to Figure 1e-j to see that discrimination cost is needed to capture the modulation across neurons.

**Figure S.3.**
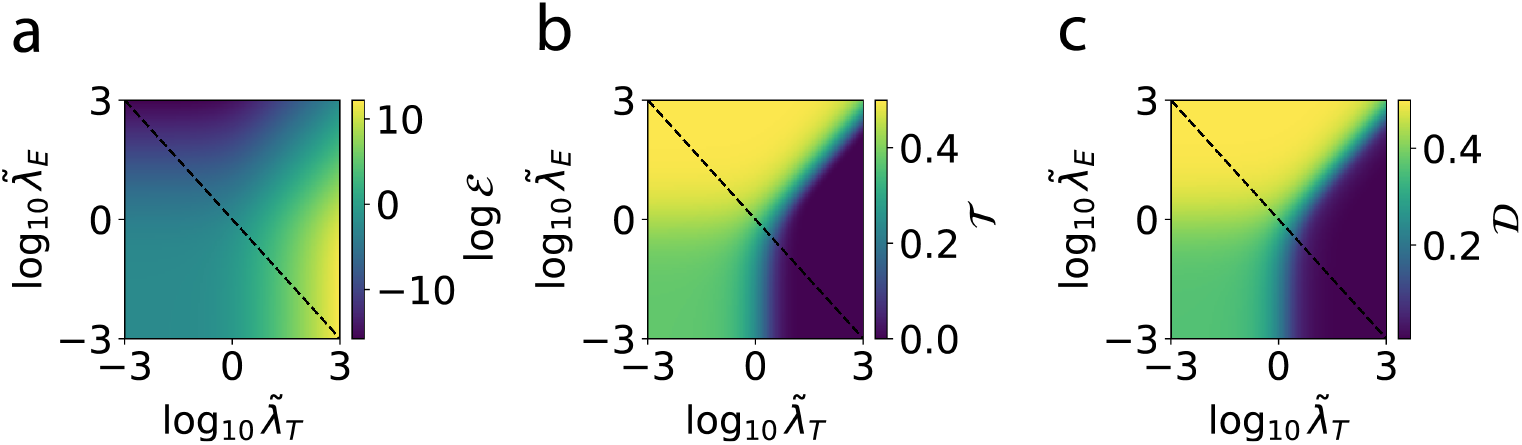
Optimizing for detection has an effect on discrimination. Optimal value for the energy cost (a), detection error (b) and discrimination error (c), as functions of the detection and energy weights (respectively the horizontal and vertical axes), rescaled as in Figure 4. Even though the discrimination term was not included in this analysis, in which we used the same cost of the simplified analytical model described in Figure 4, the optimization of the other costs end up affecting the value of the discrimination error, which closely follows the trend of the detection error. This is because changes in the overall magnitude of the responses affect both detection and discrimination errors.

**Figure S.4.**
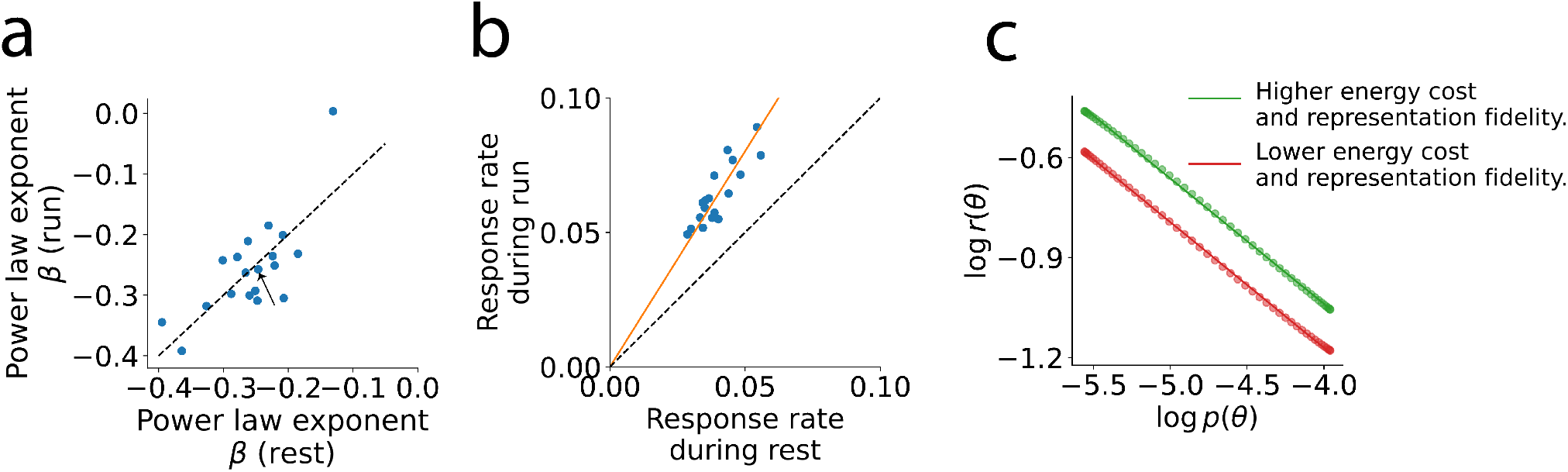
In mouse V1, locomotion increases responses without changing the power law exponent, which is reproduced by the model. a) Power law exponent for responses from the data of [1] during rest and running conditions. Dashed line: identity line. Arrow: the recording session analyzed in panel (b). b) Response rate during rest vs running states, for a specific recording session. Solid line: a linear fit with offset imposed to be zero. Dashed line: identity line. c) Two power laws with the same exponent and different prefactors obtained from the model by varying the energy and detection weights

**Figure S.5.**
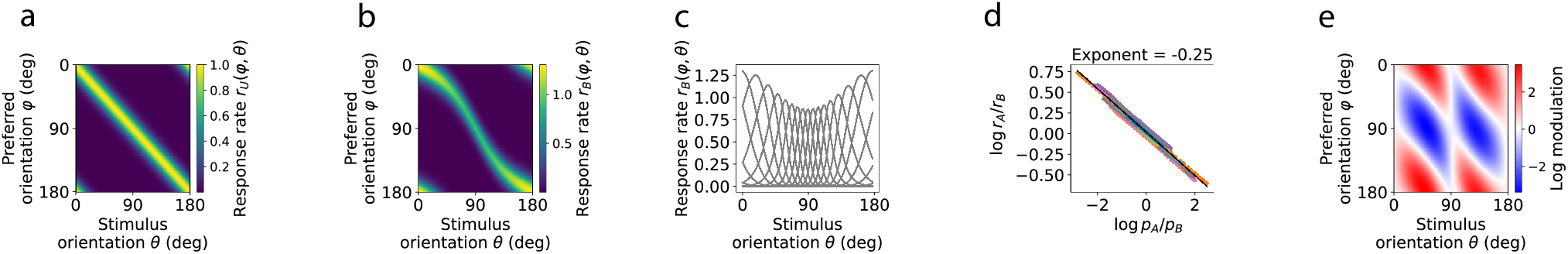
An efficient coding model on Fisher information maximization with fixed energy cost and fixed number of neurons can achieve an invariant negative power law, but with an unrealistic modulation. a-b) Predicted adapted tuning curves respectively to the uniform and a biased von Mises environment. The y-axis represents the neuron’s preferred orientation in the uniform environment. c) Same as (b), but the tuning curves are represented as different lines to appreciate the change in density and sharpness close to the adaptor. d) Ratio of average response rate vs ratio of probability of stimulus for pairs of environments. Each environment pair corresponds to a different color. e) Predicted modulation of the tuning curves obtained by taking the ratio of the biased to the uniform tuning curves.

**Figure S.6.**
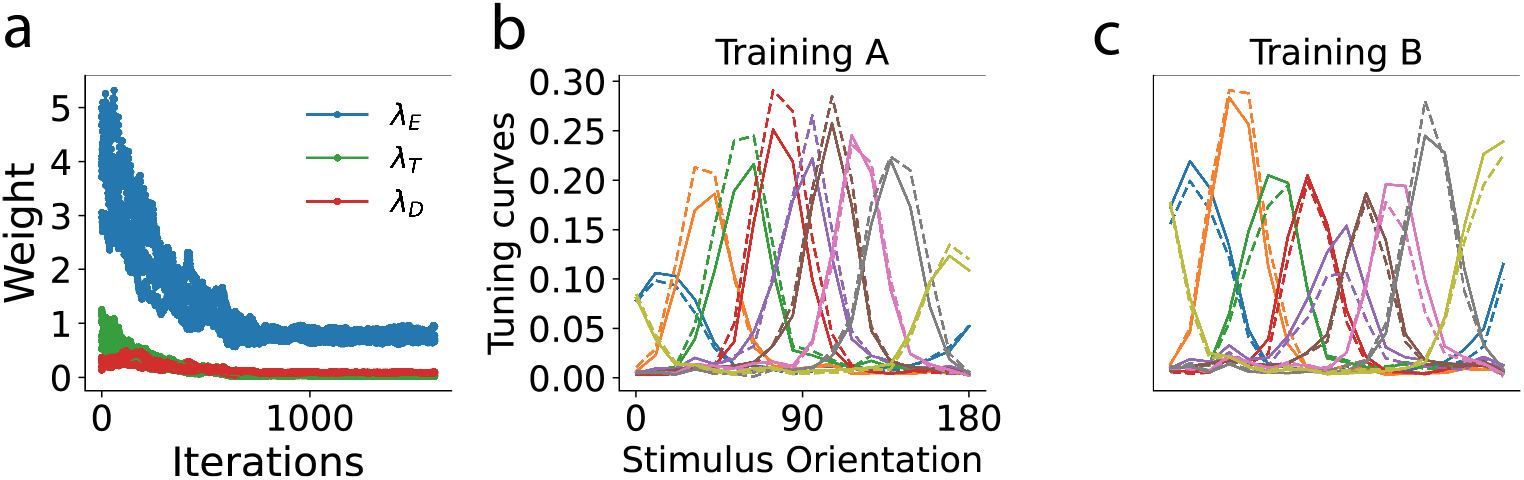
The training of the model on the data converges. a) The values of the three weights trained for a specific recording session as a function of the iterations. b-c) The tuning curves (data = dashed, converged model = solid) for the two training environments.

**Figure S.7.**
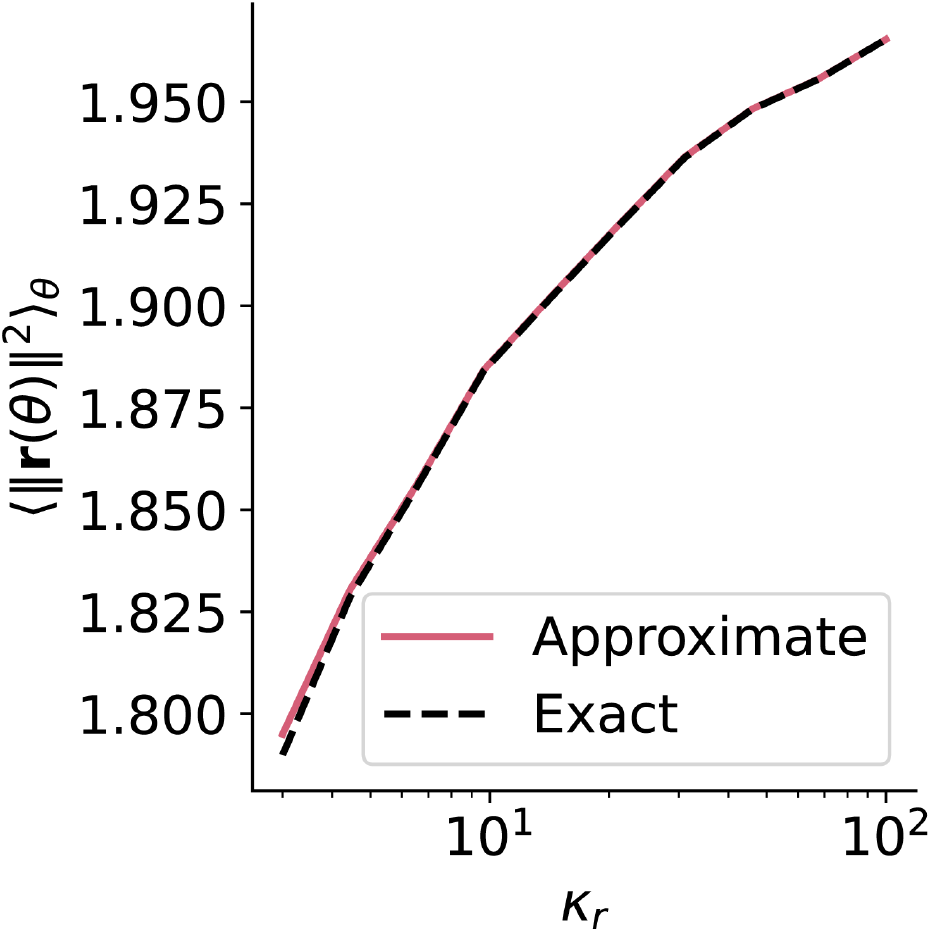
The slowly varying modulation approximation is robust for a large range of tuning sharpness. We compared the exact norm ||**r**(*θ*)||^2^ against the approximate value *a*(*θ, θ*)^2^||**r**_0_||^2^.

## Notes

### Competing Interest Statement

The authors have declared no competing interest.

### Summary of Updates

This revision substantially broadens the paper's scope. We now systematically compare our weighted-objective (energy + discrimination + detection + homeostasis) model against five prominent alternative accounts of adaptation: covariance homeostasis, product homeostasis, mutual-information maximization under limited coding capacity, reconstruction-error minimization, and a Fisher-information model with neuron-density redistribution. Only the energy-fidelity model that we introduced reproduces all the empirical signatures. Moreover, we now explain the exponent's reduction for natural stimuli with a competition between a gain-difference term and an orthogonality term, obtained with an analytic decomposition of the discrimination cost. We also implemented a Poisson-noise robustness check, a new analysis showing the exponent is unchanged across locomotion states despite amplified responses, a stronger and more rigorous analysis for the fit of the empirical responses.

